# Precocious neuronal differentiation and disrupted oxygen responses in Kabuki syndrome

**DOI:** 10.1101/484410

**Authors:** Giovanni A. Carosso, Leandros Boukas, Jonathan J. Augustin, Ha Nam Nguyen, Briana L. Winer, Gabrielle H. Cannon, Johanna D. Robertson, Li Zhang, Kasper D. Hansen, Loyal A. Goff, Hans T. Bjornsson

## Abstract

Chromatin modifiers act to coordinate gene expression changes critical to neuronal differentiation from neural stem/progenitor cells (NSPCs). Lysine-specific methyltransferase 2D *(KMT2D)* encodes a histone methyltransferase that promotes transcriptional activation, and is frequently mutated in cancers and in the majority (>70%) of patients diagnosed with the congenital, multisystem intellectual disability (ID) disorder Kabuki syndrome 1 (KS1). Critical roles for KMT2D are established in various non-neural tissues, but the effects of KMT2D loss in brain cell development have not been described. We conducted parallel studies of proliferation, differentiation, transcription, and chromatin profiling in KMT2D-deficient human and mouse models to define KMT2D-regulated functions in neurodevelopmental contexts, including adult-born hippocampal NSPCs in vivo and in vitro. We report cell-autonomous defects in proliferation, cell cycle, and survival, accompanied by early NSPC maturation in several KMT2D-deficient model systems. Transcriptional suppression in KMT2D-deficient cells indicated strong perturbation of hypoxia-responsive metabolism pathways. Functional experiments confirmed abnormalities of cellular hypoxia responses in KMT2D-deficient neural cells, and accelerated NSPC maturation in vivo. Together, our findings support a model in which loss of KMT2D function suppresses expression of oxygen-responsive gene programs important to neural progenitor maintenance, resulting in precocious neuronal differentiation in a mouse model of KS1.

**Graphical Abstract:** 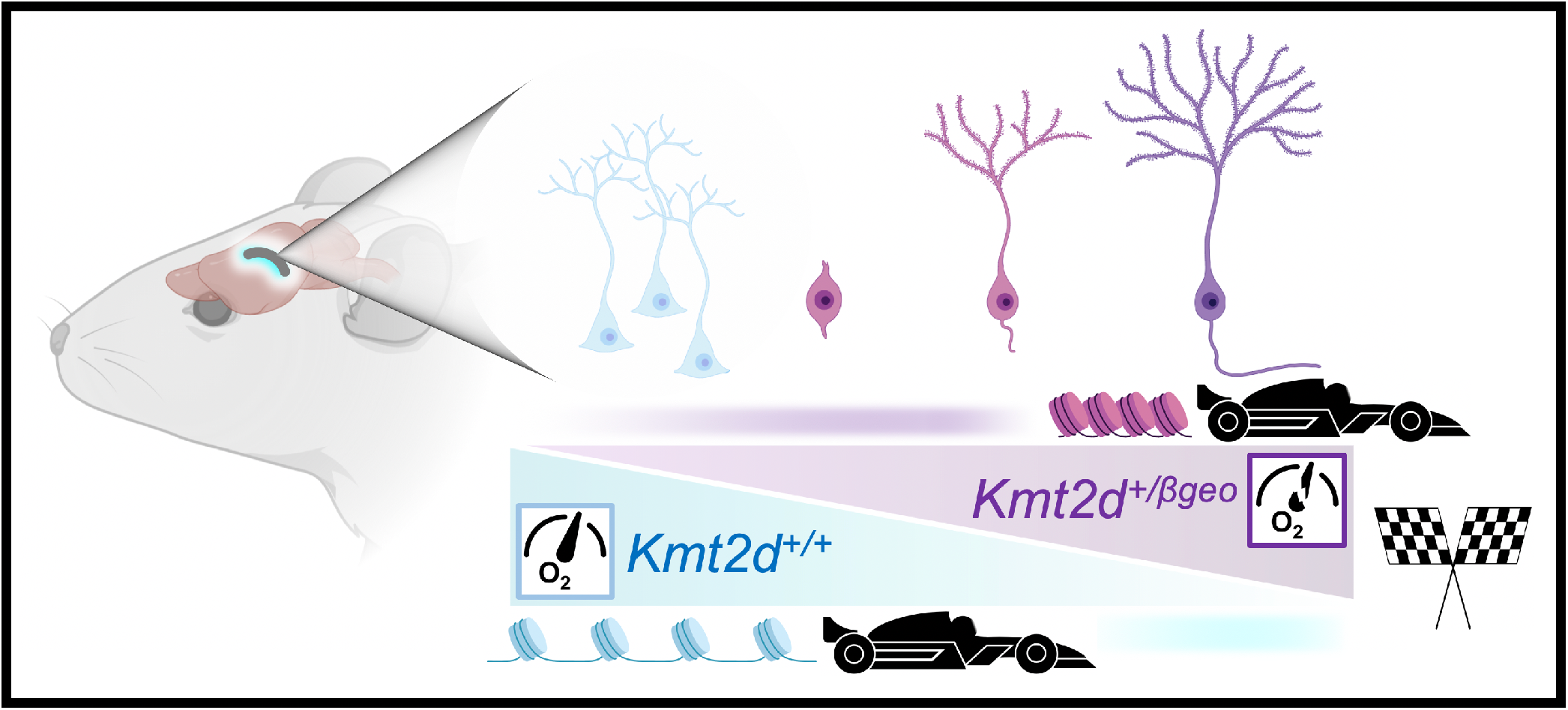

## Introduction

Trithorax group proteins promote chromatin accessibility by exerting antagonistic functions against Polycomb group transcriptional suppressors to activate gene expression (1). Fine-tuning of cell type transitions during neuronal development from NSPCs depends critically on this duality, as evidenced by severe neurodevelopmental defects caused by variants in numerous chromatin-modifying genes (2). Loss-of-function variants in genes encoding two such enzymes, KMT2D and lysine-specific demethylase 6A (KDM6A/UTX) cause the ID disorder KS (KS1 and KS2, respectively) (3, 4). Up to 74% (5) of KS cases result from mutations in *KMT2D* (KS1), encoding a major histone H3 lysine 4 (H3K4) methyltransferase which catalyzes chromatin-opening modifications at context-specific targets. Developmental requirements for KMT2D in cardiac precursors (6), B cells (7, 8), muscle and adipose (9), and epithelial tissues (10) have been linked, respectively, to *KMT2D*-associated cardiac, immunologic, and oncogenic phenotypes (11), yet the effects of KMT2D deficiency in neurodevelopment are not yet understood.

We previously described a mouse model of KS1, *Kmt2d^+/βgeo^*, demonstrating characteristic features including craniofacial abnormalities and visuospatial memory impairments, associated with decreased adult-born hippocampal NSPCs in the dentate gyrus (DG) (12). Decreased DG grey matter volume was subsequently observed in KS1 patients (13). The continual birth and integration of new neurons makes adult neurogenesis the most potent form of lifelong plasticity in the mammalian brain (14), though recent studies have disagreed on its extent in humans (15–17). During late embryonic stages, a subset of multipotent NSPCs persists in the DG (18), becoming subject to an array of intrinsic and extrinsic factors affecting NSPC maintenance, i.e. self-renewal, proliferation, and neuronal differentiation, throughout adult life. Mounting evidence tightly links metabolic rewiring (19) and hypoxic states in the DG (20, 21) to cell-intrinsic regulation of NSPC maintenance.

Here, we find that KMT2D deficiency strongly suppresses metabolic gene expression and leads to reduced proliferation, abnormal hypoxia responses, and precocious neuronal maturation in multiple KS1 model systems. Importantly, these phenotypes were validated in vivo in a KS1 mouse model, supporting a role for these abnormalities in the pathogenesis of KS1-associated ID.

## Results

### Genetic ablation of the *Kmt2d* SET methyltransferase domain disrupts proliferation and cell cycle in a cell-autonomous manner

We first selected the HT22 mouse hippocampal neuronal cell line (22) for analysis of KMT2D catalytic function in neuronal context. gDNA sequence encoding the Su(var)3-9, enhancer-of-zeste and trithorax (SET) methyltransferase domain was deleted by CRISPR-Cas9 with an upstream small guide RNA (sgRNA^up^) in exon 52, and either sgRNA^1^ (exon 54) or sgRNA^2^ (intron 54), resulting in deletions of 565 bp (*Kmt2d*^Δ1^) or 654 bp (*Kmt2d*^Δ2^), respectively, as verified by Sanger DNA sequencing, in silico translation, and PCR (Supplementary Figure 1A-B). Targeted cells were clonally expanded to establish heterozygous (*Kmt2d^+/^*^Δ^) and homozygous (*Kmt2d*^Δ/Δ^) cell lines for comparison against the parental wild-type line (*Kmt2d^+/+^*). Both biological replicate alleles, *Kmt2d*^Δ1^ and *Kmt2d*^Δ2^, were represented in present studies thus the combined data are denoted hereafter simply as *Kmt2d^+/^*^Δ^ or *Kmt2d*^Δ/Δ^. *Kmt2d* mRNA encoded within the targeted region was ∼50% decreased in *Kmt2d^+/^*^Δ^ cells and absent in *Kmt2d*^Δ/Δ^ cells, while *Kmt2d* mRNA from exons upstream of the deletion site was unaffected (Supplementary Figure 1C). Immunofluorescence against KMT2D, detecting a peptide sequence upstream of deletions (Supplementary Figure 1D), demonstrated distinctly nuclear KMT2D distribution in *Kmt2d^+/+^* cells but more diffuse distribution in *Kmt2d^+/^*^Δ^ and *Kmt2d*^Δ/Δ^ cells, while we observed uniformly nuclear expression of a neuronal nuclear marker, RNA binding protein fox-1 homolog 3 (RBFOX3), independent of genotype (Figure 1A).

**Figure 1.**
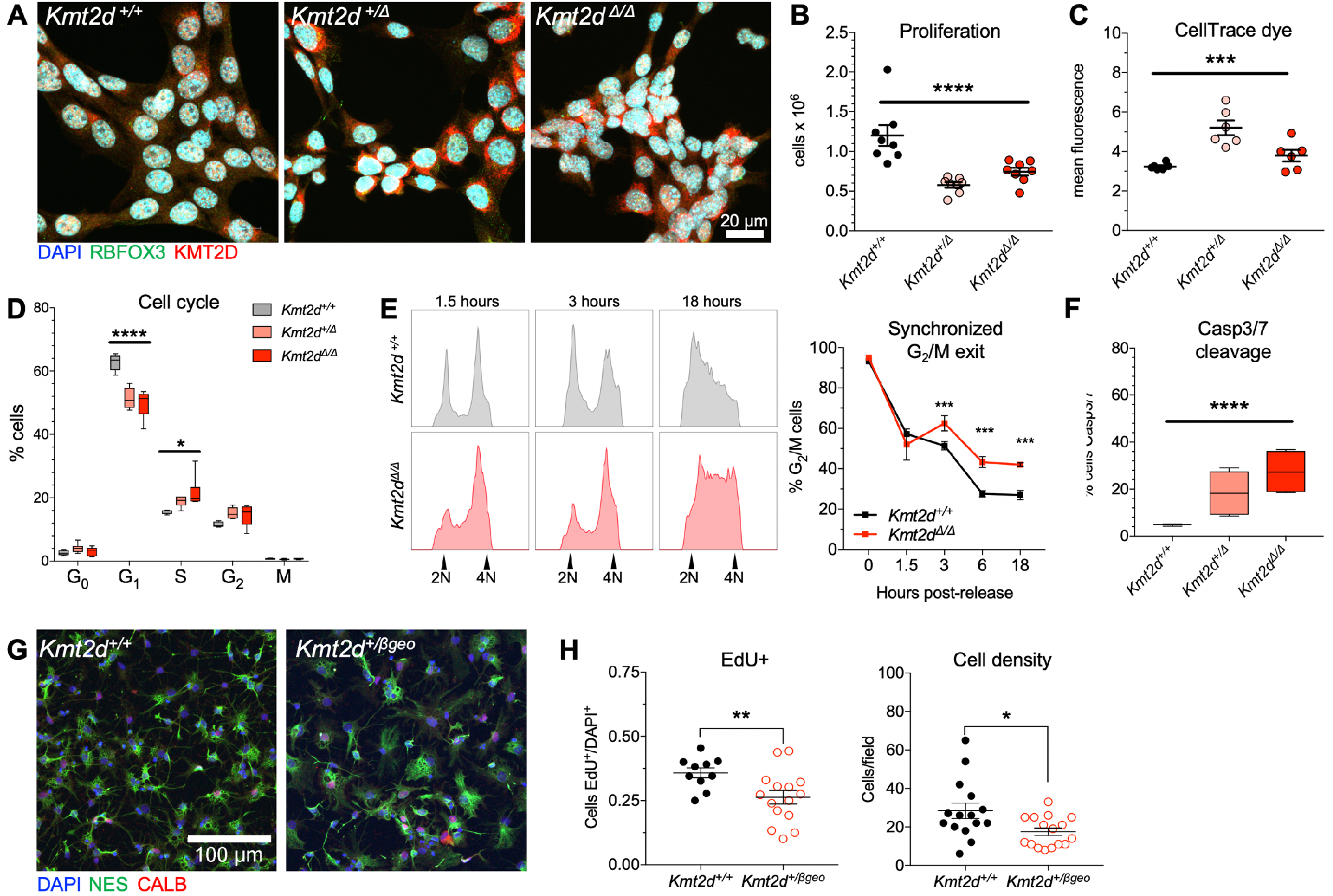
Genetic ablation of the *Kmt2d* SET methyltransferase domain disrupts proliferation and cell cycle in a cell-autonomous manner. (**A**) Representative immunostaining against KMT2D and RBFOX3 in *Kmt2d^+/+^*, *Kmt2d^+/Δ^,* and *Kmt2d^Δ/Δ^* HT22 cells. (**B**) Decreased proliferation in *Kmt2d*-inactivated cells quantified 72 hours after equal density plating. One-way ANOVA. (**C**) Generational tracking reveals fewer cell divisions, i.e. reduced dye dilution, of CellTrace Violet in *Kmt2d^+/Δ^* and *Kmt2d^Δ/Δ^* cells at 72 hours. One-way ANOVA. (**D**) Flow cytometric quantification of cell cycle phases using KI67 and DAPI fluorescence. One-way ANOVA for each cycle phase, independently. (**E**) *Kmt2d^+/+^* and *Kmt2d^Δ/Δ^* cells synchronized and released for analysis of G_2_/M exit, by DNA content, up to 18 hours after release, and quantification of cells in G_2_/M (technical triplicates per time point). Bars indicate mean ± SEM. Two-way ANOVA (P<0.0001) with post hoc multiple comparisons. (**F**) Flow cytometric quantification of early apoptotic cells by caspase-3/7 fluorescence. One-way ANOVA. (**G**) Confocal images of nestin (NES) and calbindin (CALB) expressing primary hippocampal NSPCs from *Kmt2d^+/+^* and *Kmt2d^+/βgeo^* mice, and (**H**) quantified proliferation. Student’s t-test. Bars indicate mean ± SEM. Boxes indicate mean ± interquartile range; whiskers indicate minima and maxima. (*p<0.05, **p<0.01, ***p<0.001, ****p<0.0001). Scale bars 20 μm (**A**) or 100 μm (**G**).

Proliferation analysis after equal-density plating revealed cell densities ∼52% lower in *Kmt2d*^+/Δ^ cells and ∼39% lower in *Kmt2d*^Δ/Δ^ cells, compared to wild-type (Figure 1B). This defect was supported by dye-based generational tracking, detecting modestly reduced dilution of a fluorescent tracer, i.e. fewer cell divisions, in *Kmt2d*^+/Δ^ and *Kmt2d*^Δ/Δ^ daughter cells compared to wild-type (Figure 1C, Supplementary Figure 1E), while initial dye uptake in parental cells was genotype-independent. Flow cytometric analysis of cell cycle occupancy, using marker of proliferation Ki-67 (KI67) and a DNA label, revealed that *Kmt2d*^+/Δ^ cells and *Kmt2d*^Δ/Δ^ cells were enriched for S and G_2_ phase, compared to wild-type (Figure 1D, Supplementary Figure 1F). To characterize temporal dynamics of cell cycle progression, we synchronized cells in G_2_/M phase and analyzed DNA content at timepoints after release (Figure 1E). Wild-type cells exited G_2_/M phase at higher rates than *Kmt2d*^Δ/Δ^ cells, at 3 hours and up to 18 hours after release. Cell death was profiled by flow cytometric detection of caspase-3/7 substrate cleavage to distinguish early apoptotic cells. Compared to wild-types, apoptotic cell proportions were greater in both *Kmt2d^+/^*^Δ^ cells (∼287%) and *Kmt2d*^Δ/Δ^ cells (∼478%) (Figure 1F).

To examine proliferation in primary hippocampal progenitors, we isolated NSPCs from micro-dissected DG of *Kmt2d^+/βgeo^* mice and wild-type littermates. NSPCs exhibited characteristic expression of NSPC marker nestin (NES), with a minority of cells expressing mature neuron marker calbindin (CALB) (Figure 1G). Cells were plated at equal density and pulsed with cell division marker 5-ethynyl-2′-deoxyuridine (EdU), then quantified by confocal microscopy. Compared to wild-type, *Kmt2d^+/βgeo^* NSPCs demonstrated lower proliferation rates as measured by EdU incorporation and cell density (Figure 1H).

Findings of proliferation defects, G_2_/M cell cycle delay, and increased apoptosis in hippocampal cells bearing *Kmt2d* inactivation by SET domain deletion, together with proliferation defects in primary *Kmt2d^+/βgeo^* hippocampal NSPCs, support a cell-intrinsic role for KMT2D activity in neurodevelopmental contexts.

### Suppressed transcription of KMT2D-regulated hypoxia response genes upon loss of the KMT2D SET methyltransferase domain

We performed high-coverage RNA-seq comparing three *Kmt2d*^Δ/Δ^ clones against the parental *Kmt2d^+/+^* line, each in technical triplicate, followed by differential expression analysis. Libraries clustered robustly by genotype with clear separation of *Kmt2d*^Δ/Δ^ cells from *Kmt2d^+/+^* by Principal Component Analysis (PCA), yielding 575 significant differentially-expressed genes (DEGs) at a False Discovery Rate (FDR) of 0.05 in *Kmt2d*^Δ/Δ^ cells compared to *Kmt2d^+/+^* (Figure 2A, Supplementary Figure 2A-B, **Supplementary Table 1**). ∼76% of DEGs (436 genes) were downregulated in *Kmt2d*^Δ/Δ^ cells, including known KMT2D targets such as Krueppel-like factor 10 (*Klf10*) (12), revealing strong global transcriptional suppression from *Kmt2d* inactivation. Overrepresentation analysis (ORA) revealed significant enrichment of gene networks among *Kmt2d*^Δ/Δ^ down DEGs, including glycolysis and hypoxia-inducible factor 1A (HIF1A) signaling, while *Kmt2d*^Δ/Δ^ upregulated DEGs were enriched in fewer networks (Figure 2B).

**Figure 2.**
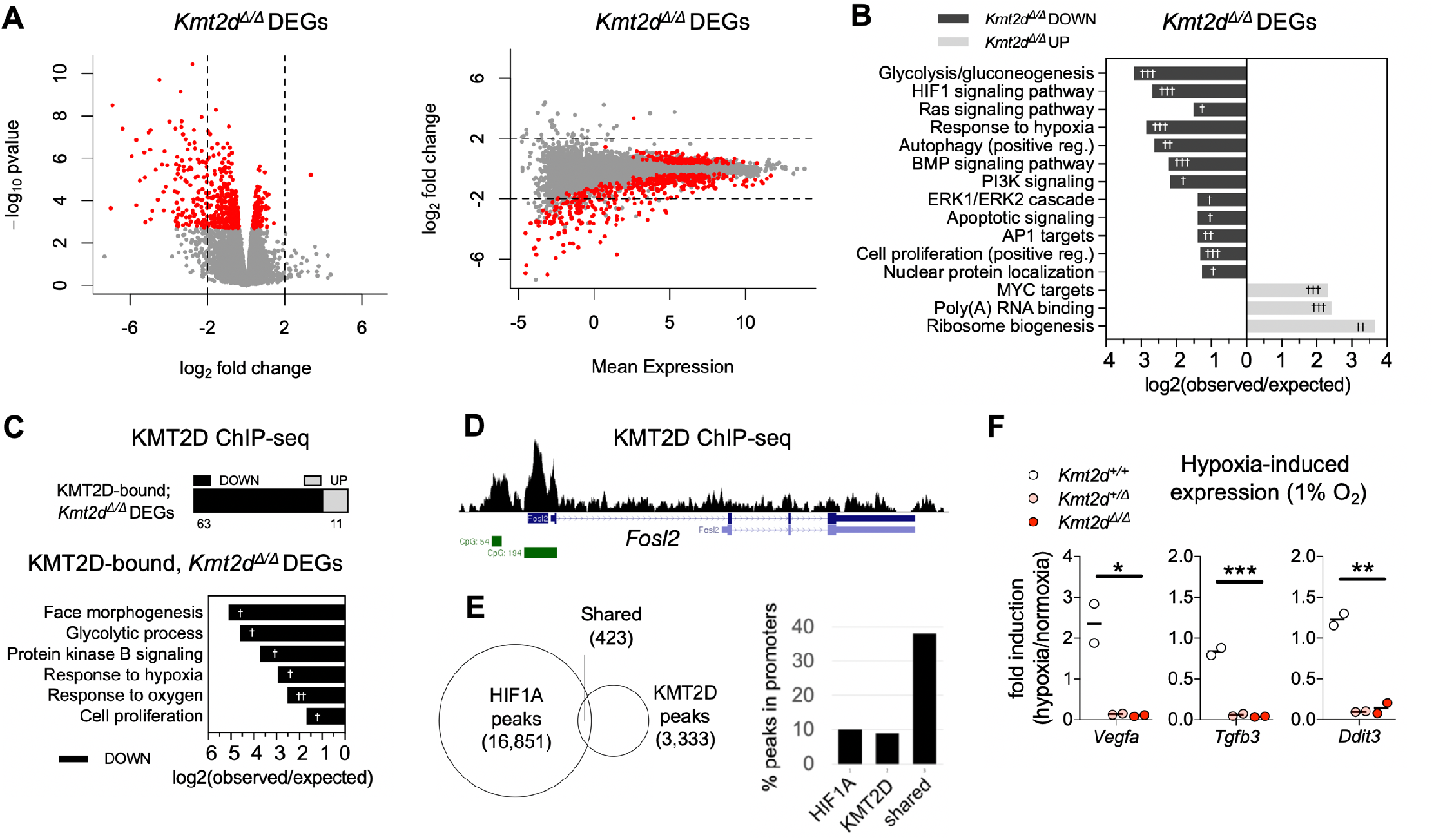
Suppressed transcription of KMT2D-regulated hypoxia response genes upon loss of the *Kmt2d* SET methyltransferase domain in neuronal cells. (**A**) Expression analysis by RNA-seq in HT22 cells reveals 575 significant differentially expressed genes (DEGs) in *Kmt2d^Δ/Δ^* clones (3 biological replicates) relative to *Kmt2d^+/+^* cells, each in technical triplicate. Fold changes in expression indicate most significant *Kmt2d^Δ/Δ^* DEGs (∼76%, red dots) are downregulated in *Kmt2d^Δ/Δ^* cells, plotted against p-value and mean expression. (**B**) Gene networks significantly enriched among down-or up-regulated *Kmt2d^Δ/Δ^* DEGs. (**C**) *Kmt2d^Δ/Δ^* DEGs which are also KMT2D-bound, as determined by ChIP-seq chromatin profiling in *Kmt2d^+/+^* HT22 cells, and gene networks significantly enriched among KMT2D-bound, *Kmt2d^Δ/Δ^* DEGs. (**D**) Representative ChIP-seq track of a KMT2D-bound, *Kmt2d^Δ/Δ^* DEG depicting KMT2D binding peaks (black), RefSeq gene annotations (blue), and CpG islands (green). (**E**) Overlapping loci of observed KMT2D-ChIP peaks in HT22 cells and HIF1A-ChIP peaks in embryonic heart (26). Overlapping KMT2D/HIF1A peak regions, compared to individually bound regions, are enriched at gene promoters. (**F**) RT-qPCR analysis of hypoxia-induced gene expression in *Kmt2d^+/+^*, *Kmt2d^+/Δ^*, and *Kmt2d^Δ/Δ^* cells, following 72 hours in normoxia (21% O_2_) or hypoxia (1% O_2_), with fold induction of target gene mRNA. 2 biological replicates per genotype, each in technical triplicate. One-way ANOVA. (*p<0.05, **p<0.01, ***p<0.001). Fisher’s Exact Test (^†^FDR<0.05, ^††^FDR<0.01, ^†††^FDR<0.001).

We reasoned that among *Kmt2d*^Δ/Δ^ DEGs, a subset of genes found to also bind KMT2D itself in wild-type cells would more likely represent direct transcriptional consequences of *Kmt2d* inactivation, whereas non-bound DEGs could reflect secondary effects. We performed chromatin immunoprecipitation followed by high-throughput sequencing (ChIP-seq) using a previously validated ChIP-grade KMT2D antibody (9) in *Kmt2d^+/+^* HT22 cells. We identified 3,756 KMT2D binding peaks significantly enriched over input (**Supplementary Table 2**), of which ∼10% occur inside promoters, ∼33% (1,235 peaks) occur within 5 kb of a transcription start site (TSS±5kb), and ∼25% occur within 2 kb (Supplementary Figure 2C-F). To account for promoter and enhancer interactions (9, 10, 23), we reasoned that TSS±5kb peaks, compared to more distal peaks, are more likely to reflect KMT2D cis-regulatory functions on proximal genes, so we refer to these as KMT2D-bound genes. The 1,463 observed KMT2D-bound genes (**Supplementary Table 3**) were significantly enriched in mRNA 3’UTR binding, rho GTPase signaling, circadian clock, translation, oxidative stress, HIF1A signaling, and other pathways (Supplementary Figure 2G).

We then intersected KMT2D-bound genes with *Kmt2d*^Δ/Δ^ DEGs to reveal 74 putative direct target genes (**Supplementary Table 3**), of which ∼85% (63 genes) were downregulated (Figure 2C), including insulin-like growth factor 1 (*Igf1*), and fos-like antigen 2 (*Fosl2*). At least 20 observed KMT2D-bound, *Kmt2d*^Δ/Δ^ DEGs were previously described as KMT2D targets in other tissues (7, 24). KMT2D-bound, *Kmt2d*^Δ/Δ^ down-DEGs were most significantly enriched for pathways including face morphogenesis, glycolysis, hypoxia response, and proliferation, and surprisingly, 29 of these 63 genes are also HIF1A-regulated (25). Although craniofacial features associate with KS1, enrichment of face morphogenesis genes in HT22 cells likely reflects pleiotropic gene functions. KMT2D ChIP-seq peaks on HIF1A-regulated genes clustered at promoters and enhancers, often overlapping CpG islands in genes such as *Fosl2*, with others clustering at alternative TSSs, as in retinoic acid receptor alpha (*Rara*), or in enhancer-like peaks, as in DNA-damage-inducible transcript 4 (*Ddit4*) (Figure 2D, Supplementary Figure 2H).

A large fraction of KMT2D-bound, *Kmt2d*^Δ/Δ^ DEGs control oxygen-responsive metabolism, warranting interrogation of shared KMT2D and HIF1A binding sites. We first intersected KMT2D peaks with HIF1A peaks previously found in embryonic heart (26), finding 423 overlapped regions (Figure 2E). Like KMT2D, HIF1A showed ∼10% of peaks located inside promoters, but among shared KMT2D/HIF1A-bound peaks this fraction approached ∼40%, supporting cooperative regulatory activity (Supplementary Figure 2I). We identified 289 TSS±5kb genes, as defined above, for these overlapped KMT2D/HIF1A-bound peaks, including 8 *Kmt2d*^Δ/Δ^ DEGs (**Supplementary Table 3**).To check if KMT2D/HIF1A-bound genes generalize to other tissues we next interrogated independent gene sets having experimentally validated, hypoxia-induced HIF1A binding in the promoter (27). Of 86 validated genes, 5 were KMT2D-bound, *Kmt2d*^Δ/Δ^ down-DEGs, 23.3-fold more than expected by chance (Fisher’s Exact Test, p=4.74e-6) (**Supplementary Table 3**). Of 81 genes validated in three or more tissues, 3 were KMT2D-bound, *Kmt2d*^Δ/Δ^ down-DEGs: *Klf10*, *Rara*, and *Ddit4* (Fisher’s Exact Test, p=0.002).

Given the prevalence of oxygen response genes among *Kmt2d*^Δ/Δ^ down-DEGs and shared KMT2D/HIF1A targets, we hypothesized a positive regulatory role for KMT2D in transcriptional responses to hypoxia in HT22 cells. We subjected *Kmt2d^+/+^*, *Kmt2d^+/^*^Δ^, and *Kmt2d*^Δ/Δ^ cells to normoxia (21% O_2_) or hypoxia (1% O_2_), and measured hypoxia-induced gene expression responses. Analysis of canonical HIF1A targets, vascular endothelial growth factor A (*Vegfa*), Bcl2/adenovirus E1B 19-KD protein-interacting protein 3 (*Bnip3*), DNA-damage-inducible transcript 3 (*Ddit3*), and cyclin-dependent kinase inhibitor 1A (*Cdkn1A*), in *Kmt2d^+/+^* cells revealed robust upregulations upon hypoxic exposure; in contrast, *Kmt2d^+/^*^Δ^ and *Kmt2d*^Δ/Δ^ cell lines failed to induce these genes to comparable levels (Figure 2F, Supplementary Figure 2J). In hypoxic conditions, stabilized HIF1A undergoes nuclear translocation, i.e. activation. We therefore quantified nucleus-localized HIF1A fluorescence under normoxia (21% O_2_) and hypoxia (1% O_2_) (Supplementary Figure 2K). Unexpectedly, in normoxia, *Kmt2d*^Δ/Δ^ cells exhibited >2-fold greater HIF1A activation than *Kmt2d^+/+^* cells. Upon hypoxic exposure, HIF1A activation doubled in wild-type cells, but failed to respond in *Kmt2d^+/^*^Δ^ cells and *Kmt2d*^Δ/Δ^ cells.

Taken together, our data suggest that KMT2D plays an important role in positively regulating HIF1A-inducible, oxygen-responsive metabolic gene programs in neuronal cells.

### KS1 patient-derived cells recapitulate KMT2D-associated defects in proliferation and cell cycle

We reprogrammed skin biopsy fibroblasts to generate induced pluripotent stem cells (iPSCs) from a previously described female KS1 patient bearing a heterozygous nonsense *KMT2D* mutation (c.7903C>T:p.R2635*) with characteristic facial features, congenital heart disease, and visuospatial memory impairments (28). We selected KS1 iPSCs (KS1-1) bearing normal 46, XX karyotype (Supplementary Figure 3A) and characteristic morphology (Figure 3A) for comparison against previously described iPSC lines from unrelated healthy controls (C1-2, C3-1) (29). *KMT2D* mRNA quantification in KS1 iPSCs confirmed decreased message compared to controls, as expected due to haploinsufficiency (Supplementary Figure 3B-C). Quantification after EdU pulse demonstrated lower proliferation rates (∼25%) in KS1 iPSCs compared to controls (Figure 3B), accompanied by a shift in cell cycle occupancy (Figure 3C, Supplementary Figure 3D) favoring G_2_/M phase (24% more cells).

**Figure 3.**
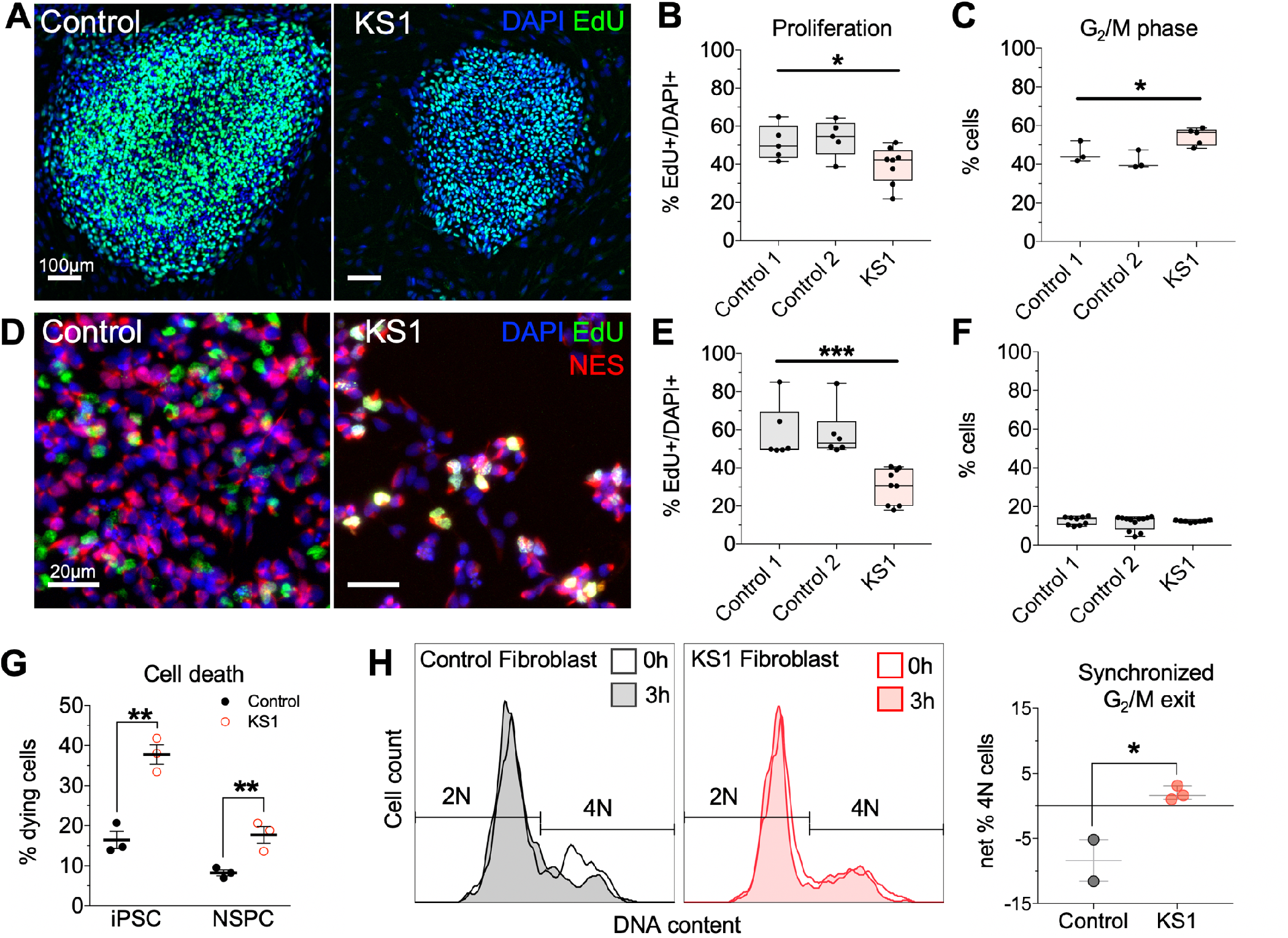
KS1 patient-derived cells recapitulate KMT2D-associated defects in proliferation and cell cycle. (**A**) Representative immunostaining of iPSCs derived from a *KMT2D^+/-^* KS1 patient (c.7903C>T:p.R2635*) and healthy controls. (**B**) Proliferating cells were pulsed with EdU for 30 minutes and quantified by flow cytometry. One-way ANOVA. (**C**) Cell cycle analysis in iPSCs, discriminating 2N and 4N DNA content (G_1_/G_0_ and G_2_/M, respectively) by flow cytometry using DAPI fluorescence. One-way ANOVA. (**D**) Representative immunostaining of NES-expressing NSPCs induced from iPSCs of KS1 patient and controls. (**E**) EdU pulse assay quantified by flow cytometry. One-way ANOVA. (**F**) Cell cycle analysis in NSPCs. One-way ANOVA. (**G**) Quantification of dying cells by flow cytometric scatter profiles in KS1 patient and control cells. Student’s t-test. (**H**) Synchronized G_2_/M exit analysis by flow cytometry in fibroblasts from KS1 patients (KS1-1, KS1-2, KS1-3) and healthy controls (Controls 3 and 4), in triplicate per cell line. Cells were enriched for G_2_/M phase using nocodazole and analyzed by DAPI fluorescence to quantify G_2_/M phase cell fractions at 0 and 3 hours after release. Student’s t-test. Bars indicate mean ± SEM. Boxes indicate mean ± interquartile range; whiskers indicate minima and maxima. (*p<0.05, **p<0.01, ***p<0.001). Scale bars 100 μm (**A**) or 20 μm (**D**).

We next generated NES-expressing NSPCs through parallel differentiation of KS1 and control iPSCs, using an established small molecule inhibition protocol (30). RT-qPCR confirmed decreased *KMT2D* in KS1 NSPCs (Supplementary Figure 3E), and cells displayed normal morphology independent of genotype (Figure 3D, Supplementary Figure 3F). EdU incorporation rates revealed KS1 NSPCs had a marked proliferation defect (∼47% reduced, Figure 3E) and fewer mitotic divisions (Supplementary Figure 3G). KS1 NSPCs did not display a cell cycle defect (Figure 3F, Supplementary Figure 3H), suggesting either cell-type dependence or loss of this phenotype during in vitro differentiation. Flow cytometry indicated higher proportions of dying cells in KS1 samples compared to controls among both iPSCs (∼130%) and NSPCs (∼115%) (Figure 3G, Supplementary Figure 3I-J).

To determine whether G_2_/M bias, seen in KS1 iPSCs, occurred in unmanipulated primary cells from additional KS1 patients, we analyzed fibroblasts from three molecularly confirmed KS1 patients (KS1-1, KS1-2, KS1-3) and healthy controls. Fibroblasts were synchronized in G_2_/M phase followed by flow cytometric analysis of DNA content. At 3 hours post-release, control cells had exited G_2_/M phase, in contrast to KS1 cells which remained in G_2_/M (Figure 3H). Thus, delayed G_2_/M exit was consistent in primary, non-reprogrammed cells from three KS1 patients.

### Transcriptional suppression of metabolic genes in cycling cells, and precocious neuronal differentiation in KS1 patient-derived NSPCs

To interrogate transcriptional consequences of *KMT2D* loss in the context of neuronal differentiation, we performed single-cell RNA sequencing (scRNA-seq) in iPSCs and NSPCs from the KS1 patient and controls (Supplementary Figure 4A). By inspecting expression of cell-type markers, we confirmed that libraries segregated into clusters reflecting distinct cell identities of the expected lineages (Supplementary Figure 4B-D).

**Figure 4.**
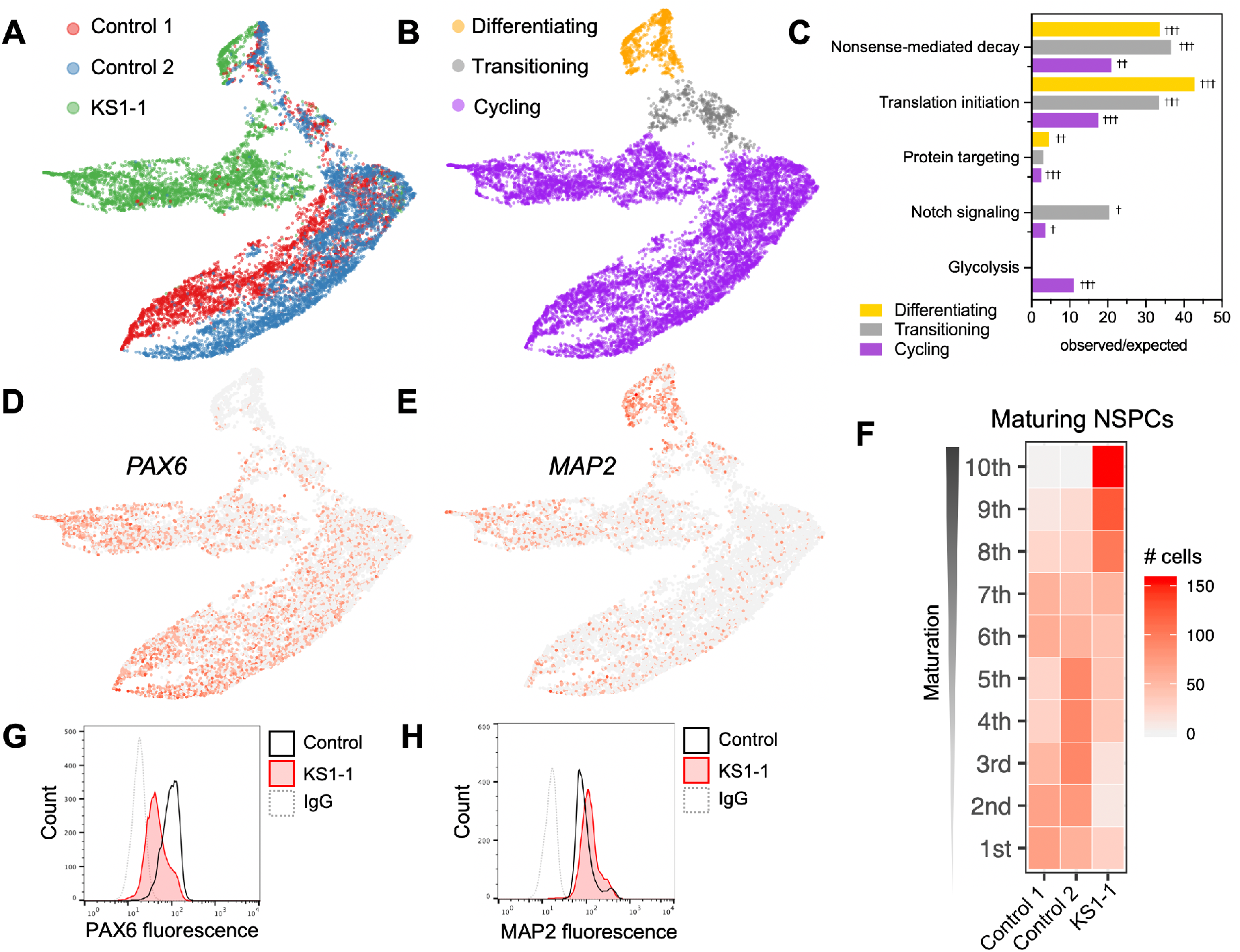
Transcriptional suppression of metabolic genes in cycling cells, and precocious neuronal differentiation in KS1 patient-derived NSPCs. (**A**) Single-cell RNA-seq profiling in patient and healthy control iPSC-derived NSPCs (∼5,000 cells per patient), with Uniform Manifold Approximation Projection (UMAP) to visualize gene expression differences between cells. (**B**) NSPCs partitioned by maturation stage as defined by stage-specific marker expression, and (**C**) enriched gene networks, analyzed exclusively among DEGs for each NSPC subset (cycling, transitioning, and differentiating). (**D-E**) Representative UMAPs annotated by relative expression intensities of NSPC markers, revealing the maturation trajectory from early NSPCs (*PAX6^+^*) to differentiating NSPCs (*MAP2^+^*). (**F**) Heatmap comparing density of NSPCs along the maturation trajectory, defined by binned marker expression from earliest (1^st^) to most differentiated (10^th^) deciles, with KS1 cells disproportionately occupying the most mature bins. (**G-H**) Protein-level experimental validation of marker expression differences by flow cytometry in NSPCs from KS1 patient and controls, plotting fluorescence intensities of PAX6 and MAP2. Fisher’s Exact Test (^†^FDR<0.05, ^††^FDR<0.01, ^†††^FDR<0.001).

First, differential expression analysis in iPSCs and NSPCs identified genes downregulated or upregulated in KS1 patient relative to healthy controls (Supplementary Figure 4E). KS1 iPSCs displayed strong transcriptional suppression among 421 DEGs, with 372 genes down and 49 genes up (**Supplementary Table 4**). NSPCs showed less directional bias, having 346 significant DEGs among which 147 genes were down and 199 genes were up (**Supplementary Table 5**). Intersection of KS1 iPSC and NSPC DEG lists showed that 40 genes were shared down and 10 genes were shared up (Supplementary Figure 4F-G, **Supplementary Table 6**). Shared down genes included glycolysis genes, aldehyde dehydrogenase 7 family member A1 (*ALDH7A1*), enolase 1 (*ENO1*), and triosephosphate isomerase 1 (*TPI1*), as well as factors important to stem cell maintenance including proliferation-associated protein 2G4 (*PA2G4*) and protein lin-28 homolog A (*LIN28A*). As in *Kmt2d*^Δ/Δ^ HT22 cells, downregulated genes in KS1 patient iPSCs and NSPCs were significantly enriched for HIF1A direct targets, genes containing the hypoxia-responsive element (HRE) 5’-RCGTG-3’ motif, and known hypoxia response genes (Supplementary Figure 4H).

We next focused on NSPCs from KS1 and controls to interrogate transcriptional effects during neuronal differentiation. We used Uniform Manifold Approximation and Projection (UMAP) to visualize single cells in a manner that displays high-dimensionality data while preserving both local and global relationships (31). Control NSPCs were tightly clustered, indicating similar expression profiles, in contrast to a distinct separation of KS1 cells which gradually lessens in a subset (top) of cells that more closely resemble controls (Figure 4A). We then partitioned single-cell libraries into developmentally informative subsets as follows. First, we verified that differences in cell cycle phase composition do not account for KS1-associated differential gene expression in NSPCs (Supplementary Figure 5A, **Supplementary Table 7**). Next, we partitioned cells by stage-specific marker expression to define a differentiation trajectory consisting of early or “cycling” NSPCs, “transitioning” NSPCs, and “differentiating” NSPCs (Figure 4B). Cycling cells comprised the majority of NSPCs analyzed and exhibited the greatest KS1-associated expression differences, while expression profiles of transitioning and differentiating NSPCs show gradual convergence of gene expression. We analyzed DEGs exclusively within cycling, transitioning, and differentiating NSPC subsets to determine if particular gene networks drive transcriptional differences in a stage-specific manner (Figure 4C, **Supplementary Table 7**). KS1 DEGs in transitioning NSPCs, and to a lesser extent cycling NSPCs, showed enrichment of genes comprising the Notch signaling pathway including delta-like protein 3 (*DLL3*), protein jagged-1 (*JAG1*), transcription factor HES-5 (*HES5*), and cyclin D1 (*CCND1*). Cycling NSPCs had DEGs enriched in glycolysis pathways.

**Figure 5.**
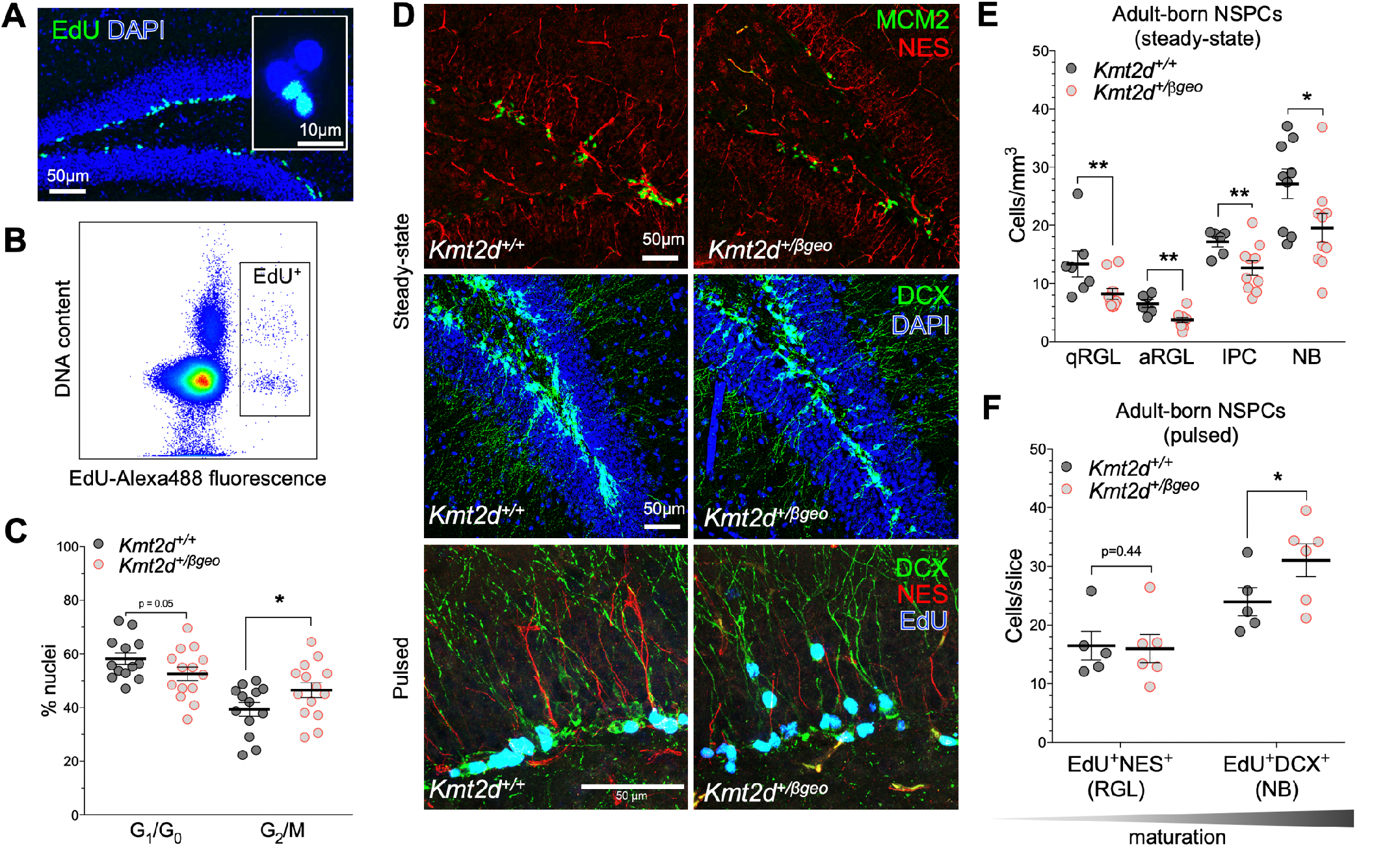
In vivo defects of neurogenesis and NSPC differentiation in a *Kmt2d^+/βgeo^* mouse model of KS1. (**A**) Immunostaining images of dividing (EdU-pulsed) dentate gyrus (DG) NSPCs, and nuclei purified from micro-dissected DG by fluorescence-activated cell sorting (FACS) (**B**) of labeled nuclei. (**C**) Cell cycle analysis in purified EdU^+^ DG nuclei from *Kmt2d^+/+^* and *Kmt2d^+/βgeo^* mice sampled 16 hours post-pulse, using DAPI fluorescence (13-14 mice per genotype, 200-500 nuclei per mouse). (**D**) Representative confocal immunostaining of neurogenesis markers in the DG of adult *Kmt2d^+/+^* and *Kmt2d^+/βgeo^* mice at steady-state (6-10 mice per genotype, 7-10 z-stack images per mouse) or after EdU pulse (5-6 mice per genotype, 10 z-stack images per mouse). NES^+^ radial glia-like (RGL) NSPCs, in either quiescent (MCM2^-^) or activated (MCM2^+^) states (qRGL and aRGL, respectively), MCM2^+^NES^-^ intermediate progenitor cells (IPCs), and DCX^+^ neuroblasts (NB) were quantified. (**E-F**) Quantification of stage-specific NSPC densities (qRGL, aRGL, IPC, and NB) in adult *Kmt2d^+/+^* and *Kmt2d^β/geo^* mice at steady-state (**E**) or after EdU pulse-chase (2 weeks) to birthdate differentiating NSPCs (**F**). Bars indicate mean ± SEM. Student’s t-test (*p<0.05, **p<0.01, ***p<0.001). Scale bars 50 μm, unless specified (**A**, inset, 10 μm).

Apart from increased rates of KS1 cell death (Figure 3G), another possible factor in the observed decrease of proliferative KS1 NSPCs (Figure 3E) could be a change in cellular differentiation, such as precocious cell maturation, resulting in depletion of cycling precursors. To explore this by scRNA-seq, we examined markers ranging from immature cells (*PAX6^+^*) to the most differentiated cells (*MAP2^+^*) (Figure 4D-E). We further restricted analysis to the transitioning and differentiating, i.e. “maturing” NSPC subset (Supplementary Figure 5B), defining a trajectory that enabled parsing of cells into binned deciles of increasing maturation (Supplementary Figure 5C-H). Quantification of cell densities revealed strong bias of KS1 NSPCs in the most matured bins relative to controls (Figure 4F), i.e. greater representation of mature NSPCs from KS1 than controls. These transcriptional signatures were corroborated experimentally at protein level, finding KS1 NSPCs had increased MAP2 fluorescence and decreased PAX6 fluorescence relative to control using flow cytometry (Figure 4G-H).

Together, these results link transcriptional suppression of metabolic gene pathways to cell-autonomous proliferation defects in *KMT2D*-deficient KS1 patient-derived stem cell models, and scRNA-seq data suggest that precocious differentiation could contribute to KS1-associated neurodevelopmental defects.

### In vivo defects of neurogenesis and NSPC differentiation in a *Kmt2d^+/βgeo^* mouse model of KS1

Finally, we asked whether proliferative defects, transcriptional suppression, and precocious differentiation phenotypes validate in vivo, using an established KS1 mouse model. *Kmt2d^+/βgeo^* mice, bearing a *Kmt2d* truncating mutation, were previously found to exhibit visuospatial memory impairments and fewer doublecortin (DCX^+^) NSPCs in the DG subgranular zone (SGZ) (12, 32), but NSPC lineage progression in *Kmt2d^+/βgeo^* mice has not been characterized.

We conducted cell cycle and RNA-seq analysis in *Kmt2d^+/βgeo^* mice, using an EdU pulse paradigm to label adult-born cells. We sampled micro-dissected DG within 1 cell cycle (16 hours) to capture the full complement of dividing NSPCs (Figure 5A), then purified EdU^+^ nuclei by fluorescence-activated cell sorting (FACS) (Figure 5B, Supplementary Figure 6A). DNA content analysis revealed enrichment of G_2_/M phase in *Kmt2d^+/βgeo^* EdU^+^ DG nuclei (Figure 5C). We next profiled transcription by RNA-seq in purified EdU^+^ DG nuclei, yielding 827 DEGs (Supplementary Figure 6B-C, **Supplementary Table 8**). The 416 down-regulated genes in *Kmt2d^+/βgeo^* nuclei were enriched for misfolded protein binding, TCA cycle, proteasome complex, oxygen response, and poly(A) RNA-binding genes. Given the observed downregulation of poly(A) RNA-binding genes, we considered the possibility that improper 3’UTR-mediated mRNA metabolism could lead to accumulation of transcripts influencing NSPC maturation. Indeed, despite little overall bias toward up or downregulation in *Kmt2d^+/βgeo^* DG nuclei, interrogating positive regulators of neuronal differentiation revealed a marked predominance of pro-neural transcripts upregulated, having only 3 genes down but 14 genes up, including copine-1 (*Cpne1*), focal adhesion kinase 1 (*Ptk2*), ras-related protein RAB11A (*Rab11A*), and retinoblastoma-associated protein 1 (*Rb1*). Interestingly, KS1 patient NSPCs also showed upregulated pro-neural genes such as nuclear receptor subfamily 2, group F, member 1 (*NR2F1*) and pro-neural transcription factor HES-1 (*HES1*), while *Kmt2d*^Δ/Δ^ HT22 cells had upregulated brain-derived neurotrophic factor (*Bdnf*) and neuron-specific microtubule component (*Tubb3/Tuj1*). Such pro-neural gene expression observed across KS1 models raises the possibility that NSPC differentiation rates could be altered in *Kmt2d^+/βgeo^* mice.

To examine NSPC lineage progression in vivo, we analyzed stage-specific cell abundances both at steady-state and after birth-dating of adult-born NSPCs by EdU pulse, comparing *Kmt2d^+/βgeo^* mice to sex- and age-matched *Kmt2d^+/+^* littermates (Figure 5D, Supplementary Figure 6D).

At steady-state, we observed significantly fewer NSPCs in *Kmt2d^+/βgeo^* mice compared to *Kmt2d^+/+^* mice at all stages (Figure 5E). The cell division marker, minichromosome maintenance complex component 2 (MCM2), distinguished NES^+^ NSPCs in the quiescent (MCM2^-^) or activated (MCM2^+^) state. Importantly, quiescent radial glia-like (qRGL, NES^+^MCM2^-^) NSPCs were ∼39% less numerous in *Kmt2d^+/βgeo^* mice, indicating a baseline paucity in the stem cell pool. Activated RGL (aRGL, NES^+^MCM2^+^) NSPCs were ∼43% less numerous, and intermediate progenitor cell (IPC, NES^-^ MCM2^+^) NSPCs were ∼26% fewer. We confirmed prior observations (12, 32) of fewer neuroblast (NB, DCX^+^) NSPCs, finding a 28% decrease in *Kmt2d^+/βgeo^* mice. By stratifying analysis along the septotemporal axis of the DG, we observed that aRGL NSPC reductions in *Kmt2d^+/βgeo^* mice were more pronounced in the septal DG than the temporal region (Supplementary Figure 6E), congruous with spatial memory defects (12) localized to the septal DG (33). Because DCX^+^ NSPCs migrate radially during maturation, we compared radial distances of DCX^+^ cell bodies from the SGZ plane and observed increased distances in *Kmt2d^+/βgeo^* mice (Supplementary Figure 6F). Finally, despite diminished NSPC populations in *Kmt2d^+/βgeo^* mice, we observed no numeric differences among mature neurons (RBFOX3^+^) (Supplementary Figure 6G), nor were gross anatomical differences seen by MRI volumetric analysis (Supplementary Figure 6H, **Supplementary Table 9**).

From these data, we then calculated a lineage progression index to approximate the expansion potential of each successive neurogenic cell type. Although *Kmt2d^+/βgeo^* mice showed fewer total NSPCs of each type at steady-state, the lineage progression index at each cell-type transition did not differ significantly (Supplementary Figure 6I), suggesting that particular cell-type transition impairments are not responsible for the adult neurogenesis defect. However, we did note substantially higher variance of RGL activation rates in *Kmt2d^+/βgeo^* mice, suggesting impaired coordination of NSPC mitotic entry (Supplementary Figure 6J).

Pulse-labeling with marker-based imaging enables precise measurement of birth dates, i.e. mitotic division, of specific cell types. To resolve temporal dynamics of NSPC differentiation in *Kmt2d^+/βgeo^* and wild-type mice, we pulsed adult mice with EdU for a period of 2 weeks, during which a subset of labeled DG cells is expected to reach a late NSPC (NB) stage, characterized by radial extension of a DCX^+^ neuronal process. In contrast, another subset of pulsed cells, bearing a NES^+^ qRGL-like process, represents NSPCs that remain in a stem-like state. Thus, by quantifying EdU-labeled cells exhibiting either a DCX^+^ neuronal process (EdU^+^DCX^+^) or a NES^+^ qRGL-like process (EdU^+^NES^+^) (Figure 5F, Supplementary Figure 7A), one can compare relative differentiation status, where a higher proportion of EdU^+^DCX^+^ cells would indicate early or precocious maturation. Indeed, though steady-state cell numbers again confirmed fewer total NES^+^ and DCX^+^ NSPCs in *Kmt2d^+/βgeo^* mice compared to wild-types, among pulsed cells the *Kmt2d^+/βgeo^* mice exhibited a significantly greater fraction of EdU^+^DCX^+^ immature neurons (Figure 5F). In other words, *Kmt2d^+/βgeo^* DG NSPCs born within the preceding 2 weeks had achieved a more advanced differentiation state than wild-type cells born in the same window.

Together, studies of adult neurogenesis dynamics in *Kmt2d^+/βgeo^* mice suggest that in vivo neurodevelopmental effects of KMT2D loss recapitulate many phenotypes observed initially in vitro using mouse HT22 cells and KS1 human-derived cells. While comparison of gene expression profiles across these KS1 models revealed few individual genes with shared dysregulation among all models, at network level we observed high enrichment of HIF1A regulatory and RNA metabolism pathways in a comparison of all DEGs in these KS1 models (Supplementary Figure 8A-D).

### Precocious differentiation and reduced hypoxia responses in *Kmt2d^+/βgeo^* primary hippocampal NSPCs

Cellular oxygen availability has previously been directly linked to maintenance and differentiation of embryonic (33) and adult DG (19) NSPCs. Primary hippocampal NSPCs of *Kmt2d^+/βgeo^* mice showed increased HIF1A activation compared to wild-type NSPCs, and both genotypes showed increased HIF1A activation upon treatment by HIF1A-stabilizing agent dimethyloxaloylglycine (DMOG) for 12 hours (Supplementary Figure 9A). We then subjected NSPCs to a standard in vitro neuronal differentiation protocol, quantifying cell marker expression between 0-8 days (Supplementary Figure 10A-B). Prior to differentiation (day 0), wild-type NSPCs expressed low levels of a mature DG neuron marker, prospero-related homeobox 1 (PROX1), while *Kmt2d^+/βgeo^* NSPCs surprisingly showed an increase (Supplementary Figure 10C). By measuring expression of a pro-neural transcription factor, achaete-scute homolog 1 (ASCL1), we observed a baseline decrease (day 0) in *Kmt2d^+/βgeo^* NSPCs compared to wild-type (Supplementary Figure 10D). In contrast, after 2 days in differentiation conditions, *Kmt2d^+/βgeo^* NSPCs responded with greater ASCL1 expression compared to wild-types, an effect sustained at 4 and 8 days. DMOG treatment increased ASCL1 levels in both genotypes, though to greater magnitude in wild-type than *Kmt2d^+/βgeo^* NSPCs. Together, these data are consistent with a link between cellular hypoxia response and neuronal differentiation in hippocampal NSPCs (20).

## Discussion

The ID disorder KS1 is caused by mutations in the histone methyltransferase *KMT2D*, but mechanistic links to neurodevelopmental and cognitive consequences in patients are not yet clear. KS1 diagnoses are typically made after childbirth, but the inherent reversibility of chromatin modifications raises the possibility that a detailed understanding of KMT2D activity in neuronal cells could identify molecular targets for postnatal interventions in KS1-associated ID.

Here, we report that KMT2D-deficient human and mouse neurodevelopment models, in vitro and in vivo, demonstrate similar patterns of transcriptional suppression, proliferative defects, and precocious cellular differentiation. These phenotypes were cell-autonomous in vitro, suggesting that 1) chromatin and gene expression studies in neurogenic cell types could yield disease-relevant KMT2D targets and 2) these cellular models provide platforms for screening of novel therapeutic strategies or targeted manipulations. We performed transcriptomic and KMT2D profiling in these models and observed systematic suppression of hypoxia response pathways, particularly among HIF1A-regulated genes that are also directly KMT2D-bound in neuronal cells. Physically overlapping KMT2D- and HIF1A-bound genomic loci were observed across tissues, ∼40% of these at promoters, raising the possibility of shared etiologies in embryonically distinct KS1-affected organ systems. Furthermore, KMT2D-deficient neuronal cells, in contrast to isogenic wild-type cells, were unable to mount characteristic hypoxia-inducible gene activation responses when exposed to low-oxygen conditions, demonstrating oxygen response defects in KS1 models.

The implication of hypoxia response defects in KS1 suggests clinical relevance of recent findings in neurodevelopmental regulation. First, the adult hippocampal NSPC niche harbors locally hypoxic, but dynamic, microenvironments and the hypoxic state positively influences NSPC survival (20, 21). Thus, compromised hypoxia responses could render cells particularly vulnerable to changes in oxygen levels as experienced by maturing NSPCs as they migrate from DG SGZ vasculature. Second, NSPC maturation is coupled to a metabolic rewiring from glycolysis in early NSPCs, to oxidative phosphorylation in maturing neurons. Zheng and colleagues (19) recently found this metabolic switch, marked by suppression of glycolytic genes, to be essential for neuronal maturation. In KS1 neural models we observed suppression of hypoxia-responsive glycolytic genes accompanied by upregulation of pro-neuronal differentiation genes, and demonstrated precocious maturation of DG NSPCs by in vivo pulsing of adult *Kmt2d^+/βgeo^* mice, as well as in vitro differentiation of *Kmt2d^+/βgeo^* primary DG NSPCs. Future studies could determine whether targeted chromatin opening at hypoxia response loci normalizes differentiation dynamics in KS1 NSPCs.

Analogous findings regarding premature activation of terminal differentiation genes, reduced proliferation, and precocious maturation in KMT2D-depleted keratinocytes were recently linked to disorganized epidermal stratification (10). Furthermore, in KMT2D-deficient cardiomyocytes, loss of H3K4me2 at KMT2D-bound hypoxia response genes associated with cell cycle and proliferative defects in heart development (6). In contrast, KMT2D deletion in B cells conferred proliferative advantage and impaired cell maturation, despite significant up-regulation of differentiation genes (8, 9). Thus, while KMT2D’s role in enhancer-mediated gene expression during differentiation is well-established (11), phenotypic manifestations appear cell type- and stage-dependent. We now extend KMT2D-associated phenotypes of transcriptional perturbance, hypoxia response, cell cycle, proliferation, and premature differentiation to neuronal contexts. Phenotypic concordance across tissues of disparate embryonic origin suggests that KMT2D targets important to KS1 phenotypes support basic cellular homeostatic functions related to housekeeping, energy production, and cell cycle progression, rather than genes with purely brain-specific function. Furthermore, we report concordant phenotypes both from nonsense *KMT2D* mutations (patient iPSCs and NSPCs), and mutations limited to the KMT2D SET domain (HT22 cells, *Kmt2d^+/βgeo^* mice), indicating that loss of either gene dosage or catalytic function of KMT2D can be pathogenic.

Present results indicate that adult hippocampal neurogenesis defects, which we previously found to associate with visuospatial memory defects in *Kmt2d^+/βgeo^* mice, are observable at all stages examined, including fewer quiescent NSPCs in the DG which could indicate either post-natal depletion or altered niche development in the embryo. Despite having fewer total NSPCs, by pulse-chase experiments we observed the *Kmt2d^+/βgeo^* NSPC population to achieve a more advanced maturation stage than that of wild-type littermates. Interestingly, adult-born NSPCs wield a disproportionately strong influence on DG circuitry and visuospatial learning during younger, but not older, neuronal maturation stages (34). This stage-dependent coupling of NSPC maturation with cognitive outcomes increases the likelihood that accelerated neuronal differentiation rates could negatively impact visuospatial memory acquisition. Furthermore, multispecies comparisons demonstrate that measured decreases in neurogenesis rates are consistent with accelerated neuronal maturation rates across the lifespan (35).

The apparent paradox of increased HIF1A activation, despite blunted hypoxia-responsive expression in *Kmt2d^+/Δ^* and *Kmt2d^Δ/Δ^* neuronal cells raises two possibilities. First, chronic HIF1A activity could result in cellular compensatory efforts to downregulate hypoxia response genes. In this case, heterochromatin environments at HIF1A-binding genes could prevent induction. Alternatively, cellular oxygen sensing could be coupled to gene expression through chromatin states in a HIF1A-independent manner. Independent studies recently discovered direct oxygen sensing by KDM6A/UTX (the H3K27 demethylase lost in KS2 patients) as well as the H3K4/H3K36 demethylase 5A (KDM5A), which controlled chromatin states and cell differentiation in a HIF1A-independent manner (36, 37). These findings link hypoxia-induced histone methylation at H3K4, H3K27, H3K9, and H3K36 directly with control of maturation in multiple cell types, further supporting the notion that KS-associated transcriptional suppression, in the adult DG context, could impact NSPC stage-dependent learning (34) via metabolic dysregulation. Hypoxia-upregulated H3K4me3 peaks (37) were enriched in HIF1A target gene promoters, where we presently observed high overlaps in KMT2D/HIF1A-bound loci. Strikingly, loss of KDM5A, whose activity opposes that of KMT2D at H3K4 sites, caused upregulation of hypoxia-responsive genes (37), i.e. an effect opposite to the present KS1-associated suppression of hypoxia response genes such as *Klf10* and *Bnip3l.* Several histone demethylases, and at least 33 chromatin modifiers in total, have been shown to impact hypoxia response genes, 11 of these associating with developmental disorders or cancers, yet KMT2D and other histone methyltransferases had not yet been implicated (38).

In summary, our findings suggest that KMT2D deficiency disrupts neurogenesis by negatively impacting NSPC maintenance functions including cell cycle, proliferation, and survival, accompanied by decreased adult NSPC numbers and precocious neuronal differentiation. Chromatin and transcriptome profiling identified KMT2D- and HIF1A-regulated gene programs suppressed across KS1 model systems, implicating previously described roles for hypoxia responses in regulating neuronal differentiation. Indeed, we functionally demonstrate KMT2D-dependent HIF1A activation and target gene induction in KS1 neural models, and diminished response to hypoxic conditions during in vitro neuronal differentiation in *Kmt2d^+/βgeo^* NSPCs. Together, these findings are consistent with an etiological model for KS1-associated developmental changes in which KMT2D loss transcriptionally suppresses oxygen response programs critical to early NSPC maintenance, favoring precocious cellular differentiation during hippocampal neurogenesis.

## Methods

Media and reagents are listed (**Supplementary Table 10**).

### Animals

The *Kmt2d^+/βgeo^* allele (Mll2Gt(RRt024)Byg) was generated by Bay Genomics (University of California) through the random insertion of a gene trap vector. *Kmt2d^+/βgeo^* mice were fully backcrossed to C57Bl/6J background (JAX) over more than 10 generations. Animals were housed in a 14-hour light/10-hour dark cycle with free access to food and water. Experiments compare age- and sex-matched littermates. Genotyping by PCR has been described (12).

### Primary hippocampal NSPCs

Female C57Bl/6J mice (JAX) were mated to *Kmt2d^+/βgeo^* males and sacrificed for embryo harvest at embryonic day 18. Micro-dissected DG from *Kmt2d^+/βgeo^* and *Kmt2d^+/+^* littermate embryos was processed for NSPC isolation and in vitro differentiation as described (39).

### Patient-derived iPSCs, NSPCs, and fibroblasts

Skin biopsy fibroblasts were cultured from molecularly confirmed KS1 patients (KS1-1, KS1-2, KS1-3). KS1-1 fibroblasts were reprogrammed using non-integrating Sendai virus vectors (CytoTune-iPS 2.0). 5 days post-induction, cells were transferred to mouse embryonic fibroblast (MEF) feeder plates in iPSC media. 21 days post-induction, high quality colonies were manually selected for propagation and karyotyping by G-banding. Generation of healthy control lines (C3-1 and C1-2) was previously described (29). Feeder MEFs from E13.5 CF-1 mice were mitotically inactivated by irradiation. iPSCs were enzymatically passaged every 4-8 days using collagenase. NSPCs were induced from iPSCs as previously described (30), briefly, by inhibiting glycogen synthase kinase 3 (GSK3), transforming growth factor β (TGF-β), *γ*-secretase, and Notch signaling pathways using small molecules CHIR99021 (4 μM), SB431542 (3 μM), and Compound E (0.1 μM), in the presence of hLIF (10 ng/ml) and ROCK inhibitor (5 μM) for 7 days. NSPCs were split with Accutase and propagated in neural induction medium on a Matrigel.

### CRISPR-Cas9 deletions in HT22 cells

HT22 mouse hippocampal cells are commercially available, but were a gift of the Goff Laboratory and maintained in HT22 media. sgRNAs targeting two loci spanning the *Kmt2d* SET domain-encoding region, with cut sites in exon 52 and either exon 54 (*Kmt2d*^Δ1^) or intron 54 (*Kmt2d*^Δ2^), were integrated into Cas9 plasmid (pSpCas9BB-2A-puro v2.0 (PX459)) and delivered to cells at 20% confluency using Lipofectamine 2000 according to manufacturer protocol. After puromycin selection, mutant cells were identified by PCR (primers listed) and clonally expanded. Following Sanger sequencing, a subset of clones appearing heterozygous by PCR, but found to bear strand invasion, were removed from analyses.

### RNA-seq in HT22 cells: library preparation

Cells were plated at equal density and sampled at 60% confluency. Total RNA was isolated from three biological replicates of *Kmt2d*^Δ/Δ^ clones and *Kmt2d*^+/+^ parental cells using Direct-Zol RNA MicroPrep, and libraries were constructed in technical triplicate using NEBNext Poly(A) Magnetic Isolation Module and NEBNext UltraII RNA Library Prep Kit for Illumina, with size selection by AMPure XP beads, according to manufacturer protocols. Library quantification and quality checks were done using KAPA Library Quantification Kit for Illumina, High Sensitivity DNA Kit on BioAnalyzer, and Qubit dsDNA HS Assay. Paired end 50 bp reads were obtained for pooled libraries using Illumina HiSeq 2500.

### RNA-seq in HT22 cells: data analysis

We first obtained a fasta file with all mouse cDNA sequences (Mus_musculus.GRCm38.cdna.all.fa.gz) from Ensembl (http://uswest.ensembl.org/Mus_musculus/Info/Index, version 91, downloaded January 2018). Then, sequencing reads were pseudoaligned to this fasta file and transcript abundances were subsequently quantified, using Salmon (40). We then used the tximport R package (41) to convert the transcript abundances into normalized gene-level counts, by setting the “countsFromAbundance” parameter equal to “lengthScaledTPM”. Next, we used the edgeR (42, 43) and limma (44) R packages to log2 transform these gene-level counts, and normalize each of the samples with the “voom” function using the effective library size (that is, the product of the library size and the normalization factors, the latter of which we computed with the “calcNormFactors” function provided in edgeR). Subsequently, we estimated the mean-variance relationship, and calculated weights for each observation. In order to account for the correlation between technical replicates of the same clone when performing the differential analysis, we fit a mixed linear model, using the function “duplicateCorrelation” from the statmod R package (45) to block on clone. The differential analysis was then performed using the limma R package. Differentially expressed genes were called with 0.05 as the cutoff for the False Discovery Rate (FDR).

When performing the principal component analysis, transcript abundances were first converted into gene-level counts using the tximport R package, with the “countsFromAbundance” parameter equal to “no”. Then, we applied a variance stabilizing transformation to these gene-level counts using the “vst” function from the DESeq2 R package (46), with the parameter “blind” set to “TRUE”, and subsequently estimated the principal components (without scaling the expression matrix) using the 1000 most variable genes.

### scRNA-seq: library preparation

NSPCs were induced in parallel from each iPSC line (KS1-1, C1-2, C3-1) under identical conditions, and passaged three times before sampling. iPSCs were detached from MEF feeders using collagenase (200 units/ml). iPSCs and NSPCs were dissociated to single-cell suspension using Accutase. Cell counts and viability were analyzed using Countess II. scRNA-seq libraries were created with Chromium Single Cell 3’ Library & Gel Bead Kit v2 (10x Genomics) according to manufacturer protocol. Targeted cell recovery for each sample was 5,000 cells. Sufficient cDNA for library construction was achieved using 20 amplification cycles for iPSC libraries and 16 cycles for NSPC libraries. Sample indexing was achieved using 11 PCR cycles for iPSC libraries and 5 cycles for NSPC libraries. scRNA-seq libraries were sequenced using Illumina NextSeq 500.

### scRNA-seq: data analysis

Sequencing output was processed through the Cell Ranger 2.1.0 preprocessing pipeline using default parameters with the exception of --expect-cells=5000 for ‘cellranger count’ and --normalize=none for ‘cellranger aggr’. Reads were quantified against hg19 using the 10x reference genome and transcriptome builds (refdata-cellranger-GRCh38-1.2.0). The aggregated raw count matrix was then used as input for the Monocle2 single-cell RNAseq framework. Differential gene expression analysis was performed on all NSPCs and iPSCs with respect to genotype (KS1 patient vs healthy control) and was performed using the Monocle2 (47) likelihood ratio test (0.1% FDR, Monocle2 LRT, Benjamini-Hochberg corrected) with ‘num_genes_expressed’ added as a nuisance parameter to both the full and reduced models. The directionality of the differential gene test was determined by calculating the mean gene expression across all KS1 patient-derived and healthy control cells respectively, evaluating the relative fold change. High-variance genes were selected as those with a positive residual to the estimated dispersion fit and a mean number of reads per cell >=0.0005. Cell cycle stage was determined by profiling cell cycle associated genes across all cells and assigning cell cycle state using the R/Bioconductor package scran (48). Dimensionality reduction and visualization was performed via UMAP (31) on the log_10_(counts + 1) of the high variance genes in the NSPC dataset. The first 10 principal components were used as input for UMAP using the default parameters of the R/CRAN package umap. Cells were assigned to clusters using Monocle2’s implementation of the louvain community detection algorithm. Learned clusters were then aggregated by hand based on marker gene expression into three clusters (“Differentiating”, “Transitioning”, “Cycling”). Differential gene expression within clusters, and between genotypes was performed as described above. The “Differentiating” cluster was then segregated, and a smooth line was fitted using a linear regression. This line was determined to represent the direction of differentiation by examination of marker genes (Supplementary Figure 5C-H). The residuals of this fit were then plotted and deciles were calculated containing equal number of cells along the axis of differentiation. The number of cells in each decile was then counted with respect to genotype.

### ChIP-seq: library preparation

*Kmt2d*^+/+^ HT22 cells were sampled at 70% confluency and processed for pull-down with ChIP-grade KMT2D antibody (Millipore Sigma) according to ENCODE guidelines. Sonicated, reverse-crosslinked chromatin served as input control. Briefly, ∼300 million cells per cell line were crosslinked in 1% formaldehyde, quenched with 0.125 M glycine, and cell lysate supernatants were collected for immediate processing or snap-frozen for storage at −80°C. Nuclei (20 million/sample) were diluted in 1 ml RIPA buffer were sonicated using Bioruptor for 6 cycles of 5 minutes (60 seconds on/30 seconds off) in ice-cold water bath. Supernatants containing sheared chromatin were pre-cleared with Protein A Dynabeads and incubated overnight at 4°C with 8 μg KMT2D antibody. ChIP DNA was recovered by Dynabead incubation (overnight at 4°C plus 6 hours at room temperature) before 6 sequential salt washes of increasing stringency, then eluted and reverse crosslinked overnight at 65°C. DNA was purified using DNA Clean and Concentrator (Zymo Research) and quantified using High Sensitivity DNA Kit on BioAnalyzer, and Qubit dsDNA HS Assay. DNA libraries were constructed using NEBNext UltraII DNA Library Prep Kit for Illumina and quantified using KAPA Library Quantification Kit for Illumina. Paired end 75 bp reads were obtained for pooled libraries using Illumina HiSeq 2500.

### ChIP-seq: data analysis

Sequencing reads were aligned to the mouse reference genome (mm10) using Bowtie2 (49). Then, duplicate reads were removed with the function MarkDuplicates from Picard (http://broadinstitute.github.io/picard/). Peaks were subsequently called using MACS2 (50), with the “keep-dup” parameter equal to “all”. After peak calling, we excluded all peaks that overlapped with blacklisted regions provided by ENCODE (51). As a quality metric, using the resulting list of peaks, we computed the fraction of reads in peaks (frip) with the “featureCounts” function in the Rsubread package (52), with the “requireBothEndsMapped” parameter equal to “TRUE”, and the “countChimericFragments” and “countMultiMappingReads” parameters equal to FALSE. We found frip to be 2.9%, which is within the typically encountered range of values for a point-source factor (53). To identify genes likely to be regulated in *cis* by KMT2D, we first obtained the coordinates of 10kb regions centered around the TSS for each gene, using the “promoters” function from the EnsDb.Mmusculus.v79 R package (54), with the “filter” parameter equal to “TxBiotypeFilter(“protein_coding”)”, and the “upstream” and “downstream” parameters both equal to 5000. Subsequently, we selected those genes whose extended promoter (+/- 5kb from the TSS) overlapped with at least one KMT2D peak, using the “findOverlaps” function in the GenomicRanges R package (55).

### Purification of EdU^+^ nuclei

Mice were given 150 mg/kg EdU by intraperitoneal injection and sampled after 16 hours. DG was micro-dissected in ice-cold PBS immediately following sacrifice by halothane inhalation. Total nuclei were purified as described (56) with addition of RNase inhibitor to all buffers. Briefly, DG was dounce-homogenized in 1 ml lysis buffer and layered above a sucrose gradient for ultracentrifugation at 28,600 RPM for 2 hours at 4°C. Nuclei were resuspended in Click-iT EdU AlexaFluor-488 with RNAse inhibitor, and incubated 30 minutes at room temperature. Samples were passed through 40 μm filter, stained with 1 μg/ml DAPI, and kept on ice before sorting. Lysates processed identically from non-EdU-injected mice served as negative controls during sorting with Beckman Coulter MoFlo Cell Sorter. Cell cycle analysis by DNA content was performed with gates discriminating 2N and 4N cells by DAPI fluorescence.

### RNA-seq: EdU^+^ nuclei

Purified EdU^+^ nuclei from 3 *Kmt2d^+/βgeo^* and 3 wild-type littermate female mice (500 nuclei pooled per genotype) were sorted into Smart-Seq 2 lysis buffer (2 µL Smart-Seq2 lysis buffer with RNase inhibitor, 1 µL oligo-dT primer, and 1 µL dNTPs), briefly spun by tabletop microcentrifuge, and snap-frozen on dry ice. Nuclei were processed according to a modified Smart-seq2 protocol (57). Briefly, lysates were thawed to 4°C, heated to 72°C for 5 minutes, and immediately placed on ice. Template-switching first-strand cDNA synthesis was performed using a 5’-biotinylated TSO oligo. cDNAs were amplified using 20 cycles of KAPA HiFi PCR and 5’-biotinylated ISPCR primer. Amplified cDNA was cleaned using 1:1 ratio of Ampure XP beads and approximately 250 pg was input to a one-quarter-sized Nextera XT tagmentation reaction. Tagmented fragments were amplified for 12 enrichment cycles and dual indexes were added to each well to uniquely label each library. Concentrations were assessed with Quant-iT PicoGreen dsDNA Reagent (Invitrogen) and samples were diluted to ∼2nM and pooled. Pooled libraries were sequenced on the Illumina HiSeq 2500 platform to a target mean depth of ∼8 x 105 bp paired-end fragments per cycle. Paired-end reads were aligned to mm10 using HISAT2 (58) with default parameters except: -p 8. Aligned reads from individual samples were quantified against a reference genome (GENCODE vM8) using cuffquant (59). Normalized expression estimates across all samples were obtained using cuffnorm with default parameters (60).

### RT-qPCR

Total RNA was isolated by RNeasy Mini and cDNA libraries were constructed with High-Capacity cDNA Reverse Transcription Kit (Applied Biosystems) according to manufacturer protocols. Experiments were performed in technical triplicate, with biological replicates as indicated. Probes were from Taqman.

### Immunostaining, confocal imaging, and processing

Coronal brain sections of 30 μm (every sixth slice) were analyzed in serial order. Briefly, adult brains were PFA-fixed by transcardial perfusion and post-fixed for 12 hours before cryoprotection by 30% sucrose in phosphate buffer. Brains were sectioned by cryostat (Leica), directly mounted to charged slides, and stored at −80°C. Antigen retrieval (DakoCytomation) was performed at 95°C for 20 minutes. Overnight incubation at 4°C in primary antibodies (**Supplementary Table 10**) preceded AlexaFluor-conjugated secondary antibody (1:500). Tiled, z-stacked images were acquired using Zeiss LSM780 FCS AxioObserver confocal microscope and Zen software (Zeiss) to encompass entire DG structure. Images were quantified using Imaris (BitPlane) by experimenters blinded to genotype. Cell counts were corrected by DG area multiplied by z-thickness, and expressed as cells/mm^3^. For pulse-label experiments, mice were injected with 150 mg/kg EdU in saline every 48 hours and sampled as above. DCX^+^ neuroblast distance from SGZ plane was measured in Fiji (NIH). Patient-derived cell imaging utilized EVOS FL Cell Imaging System with analysis in Fiji.

### FACS and analysis

Flow cytometry analysis with FACSverse and FACSsuite (BD Biosciences), and sorting by Beckman Coulter MoFlo Cell Sorter with proper gate settings and doublet discrimination (Supplementary Figure 3J, Supplementary Figure 6A). Runs of 10,000 or more cells were analyzed from technical triplicate culture wells and analyzed in FlowJo v10 (Tree Star Inc). Unstained and secondary-only samples served as control. Cells were sampled after 30-minute pulse of EdU (10 μM) using Click-iT EdU Flow Cytometry Assay (ThermoFisher Scientific). CellTrace Violet and CellEvent caspase-3/7 reagent (ThermoFisher Scientific) were used according to manufacturer protocols. For cycle synchronization, 250 ng/ml nocodazole (Sigma) was applied for 18 hours before release.

### Magnetic Resonance Imaging (MRI)

3D T2-weighted MRI (9.4T) was performed in PFA-perfused brains of *Kmt2d^+/βgeo^* (n=3) and *Kmt2d^+/+^* (n=3) female mice aged 4 months. Atlas-based, volume-corrected analysis was performed in 25 brain regions (DtiStudio).

### Statistics

For high-throughput experiments, see Methods. For cellular assays, see Figure Legends. Statistical analyses with multiple comparisons correction were done with GraphPad Prism (version 7.0b). Gene set enrichments were determined according to WebGestalt (61), or by Fisher’s Exact Test in R version 3.5.2 as indicated.

### Study Approval

All mouse experiments were performed using protocols approved by the Animal Care and Use Committee of Johns Hopkins University School of Medicine and are in accordance with NIH guidelines. Informed consent regarding KS1 patient samples was obtained according to institutional IRB and ISCRO protocols approved by JHU.

## Author contributions

GAC and HTB conceived the study; GAC and HTB wrote the manuscript; GAC, HNN, GC, JDR, LZ performed experiments; GAC, LB, JA, KDH and LG analyzed data.

## Acknowledgments

HTB is funded through an Early Independence Award from the National Institutes of Health (NIH, DP5OD017877), the Icelandic Research Fund (195835–051), and the Louma G. Foundation. Imaging was performed with NIH support (S10OD016374). Karyotype facility supported by NICHD (1U54 HD079123-01A1). FACS was performed at the Bloomberg School of Public Health. We thank Michael Sherman for assistance with image quantification and Manisha Aggarwal for MRI. Schematics created with Mark Sandusky or Biorender.com. Hongjun Song and Kai Ge provided critical reagents and advice. Hal Dietz and Gregg Semenza provided conceptual guidance.

## Data availability

High-throughput data are publicly available. RNA-seq and ChIP-seq: GEO #GSE126167. scRNA-seq: GEO #GSE126027. Scripts for scRNA-seq analysis are available at https://github.com/Jaugust7/Kabuki-Syndrome-scRNA-analysis.

## Supplementary Figures

**Supplementary Figure 1.**
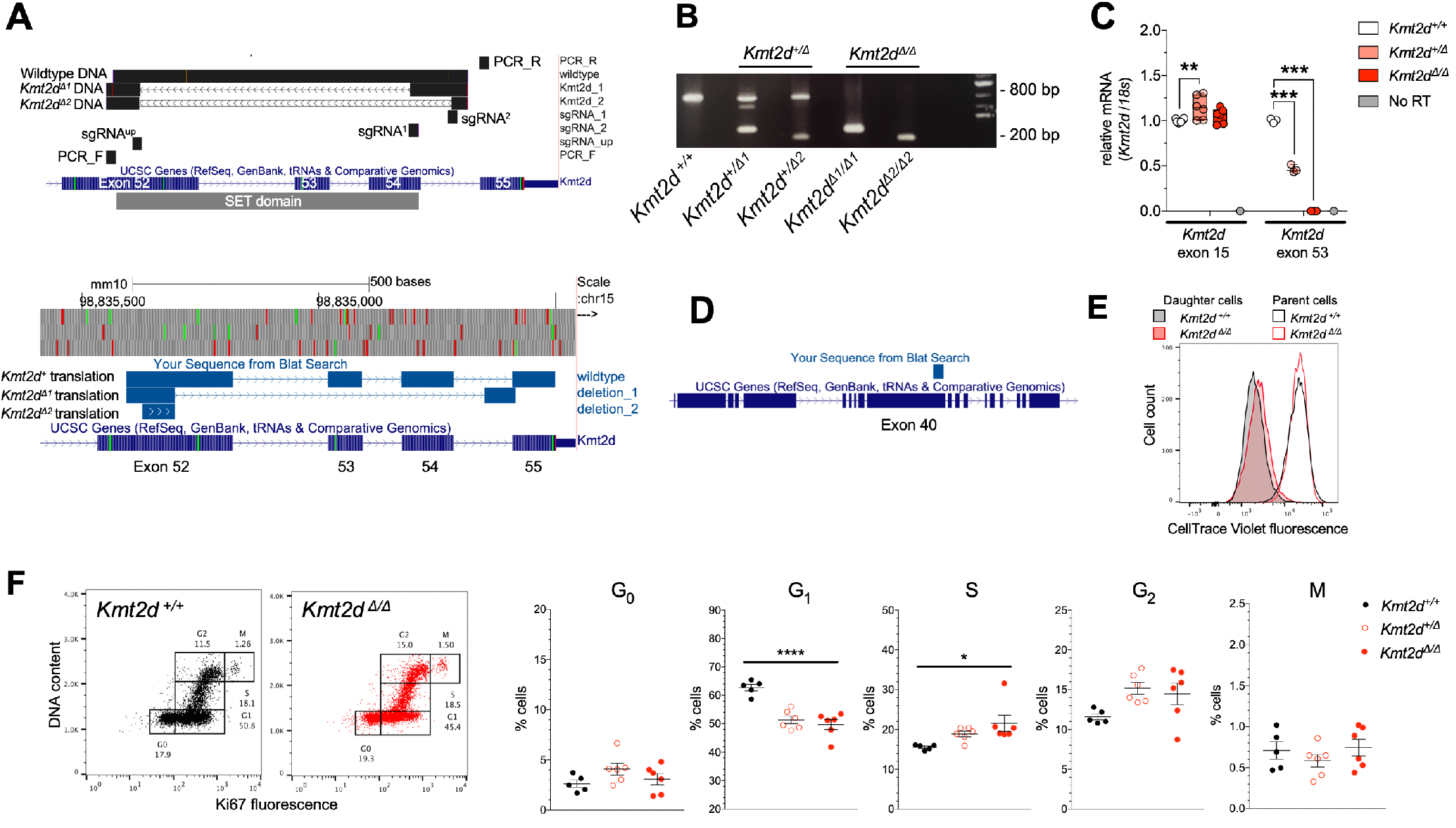
CRISPR-targeted HT22 cells. (**A**) Sanger-sequenced DNA of wild-type (*Kmt2d^+^*) and targeted (*Kmt2d^Δ^*) alleles in HT22 cells, mapped with sgRNAs and PCR primers, to *Kmt2d* locus (mm10) on chromosome 15. Mapping of Sanger-sequenced DNA after in silico translation to predict amino acid sequences illustrates premature termination codons (PTC) created in *Kmt2d^Δ1^* and *Kmt2d^Δ2^* alleles. (**B**) PCR with probes flanking sgRNA cut sites identifies experimental cell lines (*Kmt2d ^+/Δ^* and *Kmt2d^Δ/Δ^*) compared to wild-type (*Kmt2d^+/+^*). (**C**) RT-qPCR analysis of mRNA using probes spanning upstream exons (15–16) or exons within the deletion site (53–54). Two-way ANOVA with post hoc multiple comparisons. (**D**) Mapped peptide sequence of KMT2D antibody (Sigma). (**E**) Flow cytometric CellTrace fluorescence after 72 hours in HT22 cells. Increased intensity indicates less dye dilution, i.e. fewer cell divisions in mutants (left). Parental cell data confirm genotype-independent dye uptake (right) at 0 hours. (**F**) Cell cycle gating by flow cytometric analysis using Ki67 and DAPI to discriminate individual stages (G_0_, G_1_, S, G_2_, M) in *Kmt2d^+/+^* and *Kmt2d^Δ/Δ^* cells, and quantification of each cycle phase. One-way ANOVA. Bars indicate mean ± SEM. Boxes indicate mean ± interquartile range; whiskers indicate minima and maxima. (*p<0.05, **p<0.01, ***p<0.001, ****p<0.0001).

**Supplementary Figure 2.**
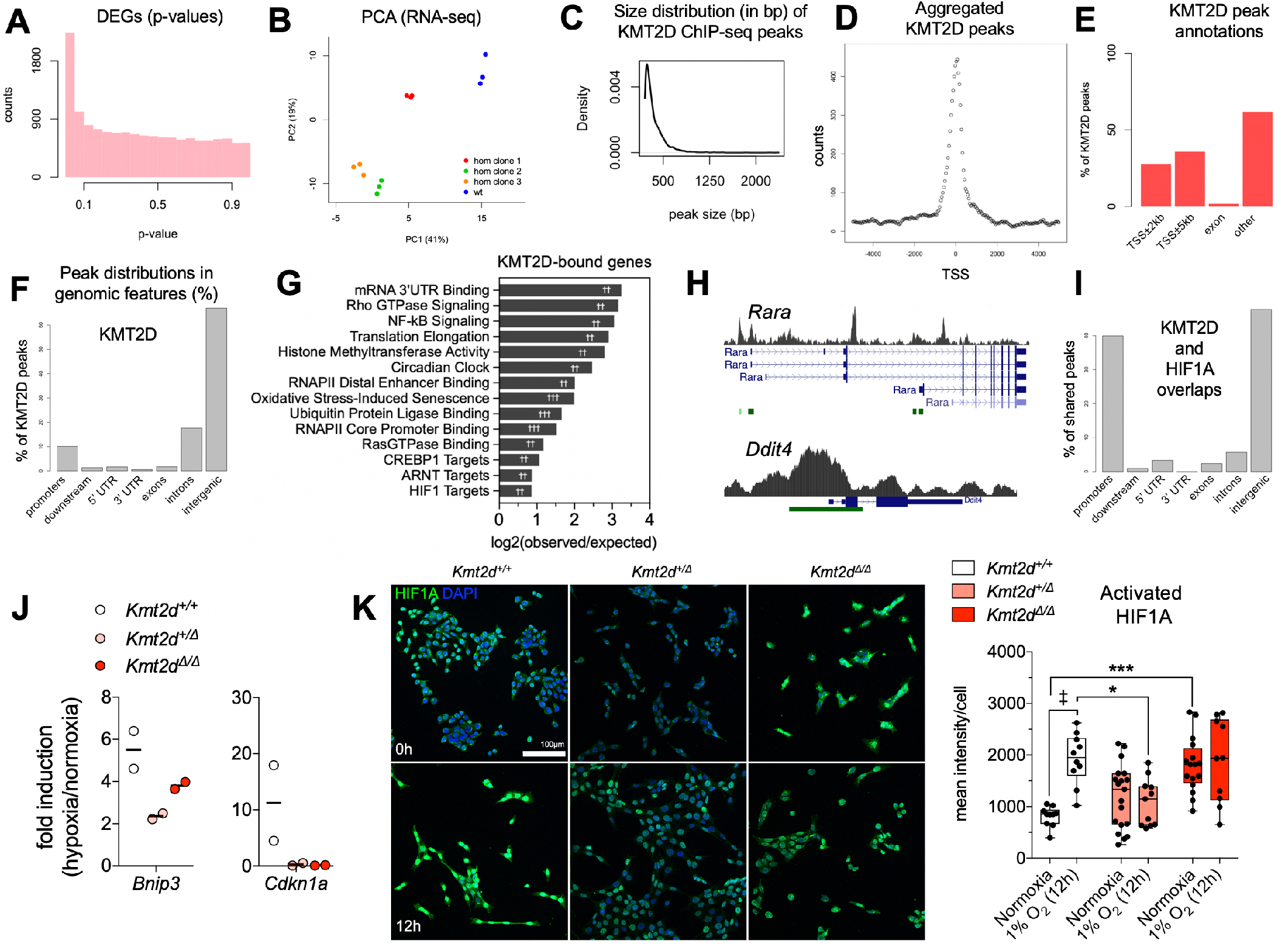
HT22 cell RNA-seq and ChIP-seq analysis. (**A**) P-value distribution in *Kmt2d^Δ/Δ^* DEGs relative to wild-type indicates a well-calibrated test. (**B**) PCA visualizing clear expression differences in wild-type and *Kmt2d^Δ/Δ^* HT22 cells. (**C**) The size distribution (in bp) of KMT2D ChIP-seq peaks. (**D**) Validation of KMT2D peak distributions about gene TSSs and (**E-F**) genomic features. (**G**) Gene networks showing highest fold change in enrichment among genes proximal to KMT2D peaks (TSS±5 kb). Fisher’s Exact Test (^†^FDR<0.05, ^††^FDR<0.01, ^†††^FDR<0.001). (**H**) KMT2D peaks clustered at alternate TSSs of *Rara* gene and enhancer-like peaks at *Ddit4* gene. (**I**) Genomic features at overlapping KMT2D and HIF1A (26) ChIP-seq peaks. (**J**) RT-qPCR analysis of hypoxia-induced gene expression in HT22 cells, upon 1% O_2_ exposure. One-way ANOVA (n.s.). (**K**) HIF1A nuclear fluorescence, i.e. activation, analysis. Representative z-stacked confocal images are shown with quantifications of nuclear HIF1A fluorescence. Two-way ANOVA with post hoc multiple comparisons (significance from wild-type, *p<0.05, **p<0.01, ***p<0.001; and from baseline, ^‡^p<0.01). Boxes indicate mean ± interquartile range; whiskers indicate minima and maxima. Scale bar 100 μm.

**Supplementary Figure 3.**
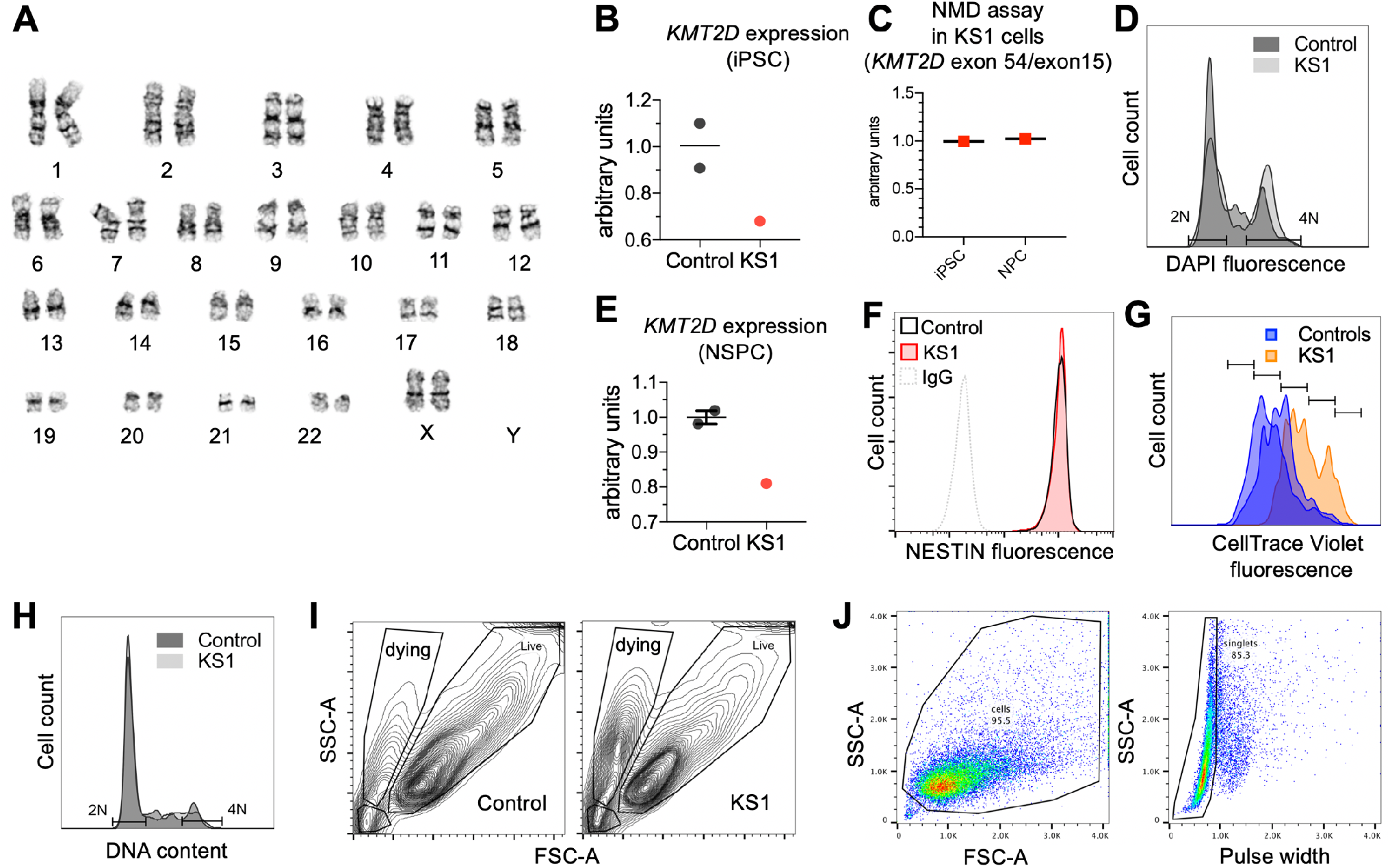
iPSC and NSPC line validations and additional phenotyping. (**A**) 46, XX normal female karyotype in KS1-1 iPSCs. (**B**) RT-qPCR analysis of *KMT2D* (exon 15) expression in KS1 iPSCs compared to two healthy control iPSC lines (C1-2 and C3-1). Dots represent average of technical triplicates per patient line. Bars indicate mean. (**C**) RT-qPCR demonstrating equivalent exonic ratios of *KMT2D* exon 15 to exon 54, measured in technical triplicate, consistent with NMD of the entire transcript. (**D**) Flow cytometric analysis of DNA content by DAPI fluorescence in iPSCs. (**E**) RT-qPCR analysis of *KMT2D* (exon 15) expression in NSPCs derived from the KS1 and control iPSC lines, measured in technical triplicate. (**F**) Flow cytometric analysis of NES fluorescence intensity in KS1 and control NSPCs. (**G**) CellTrace Violet generational tracking showing fewer divisions (i.e. higher dye intensity) in patient-derived NSPCs over 72 hours. (**H**) Flow cytometric analysis of DNA content by DAPI fluorescence in NSPCs. (**I**) Sample flow cytometric gating for detection of scatter profiles indicative of cell death-associated cellular condensation. (**J**) Representative gating of viable cells and doublet discrimination in immunofluorescence-based flow cytometric analyses of iPSCs and NSPCs.

**Supplementary Figure 4.**
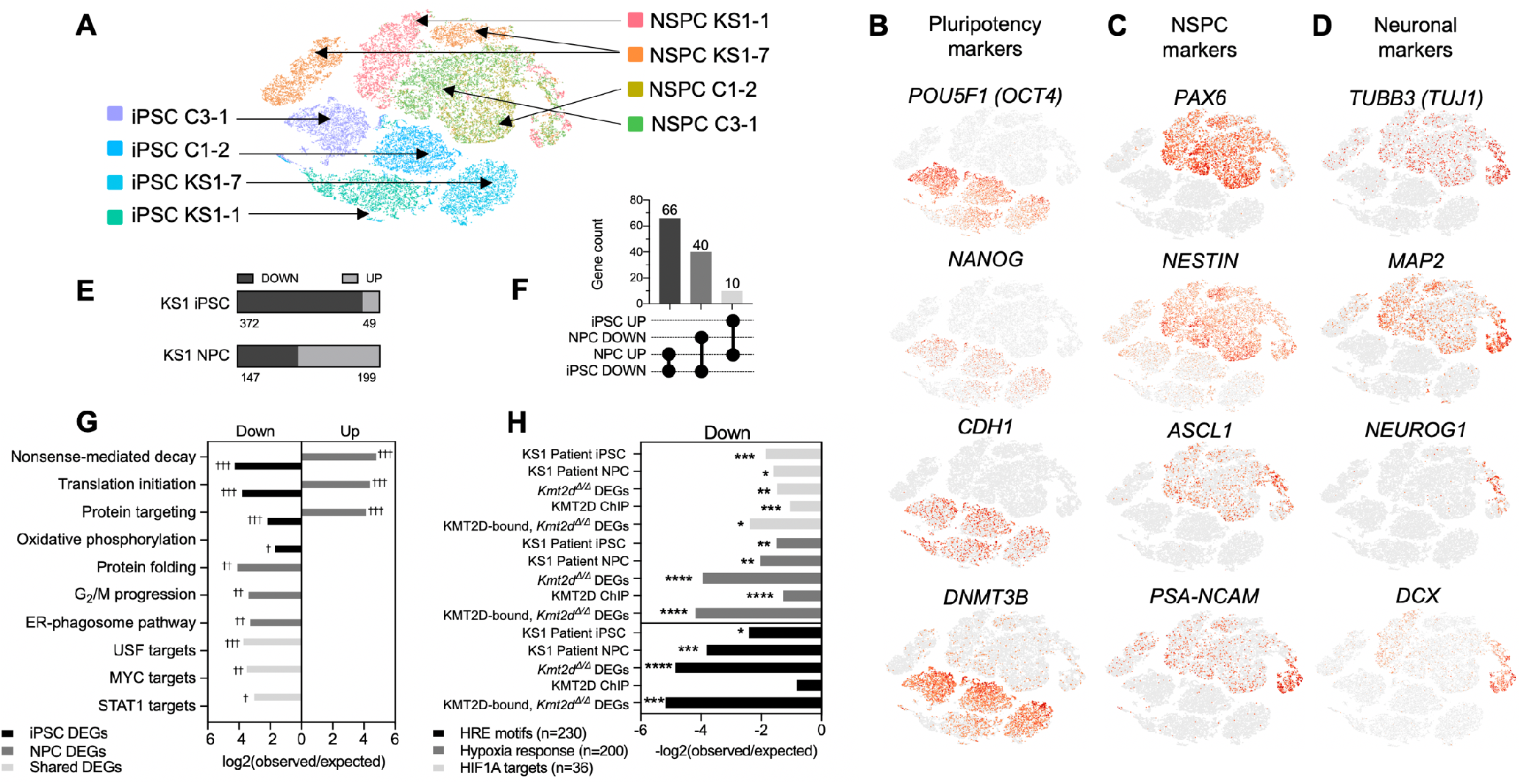
iPSC and NSPC single-cell RNA-seq analysis. (**A**) t-stochastic neighbor embedding (tSNE) representation of iPSC and NSPC libraries sequenced on 10XGenomics platform. Cell clusters colored by cell type and patient ID. iPSCs and NSPCs derived from patient K1-7 were excluded from downstream analysis due to abnormal karyotype. (**B-D**) Representative tSNE of iPSC, NSPC, and neuronal markers demonstrating expected cell identities and revealing a gradient of cell maturation. (**E**) Proportions of DEGs down- or up-regulated in KS1 patient iPSCs or NSPCs compared to respective healthy controls, (**F**) DEG lists intersected for overlaps among down-regulated and up-regulated genes, and (**G**) Gene networks most enriched among differentially expressed genes (DEGs) in KS1 patient iPSCs and NSPCs relative to respective healthy controls, and DEGs shared in both cell types. (**H**) Significant enrichments of Hypoxia Response genes, HIF1A Direct Target genes, and genes containing the Hypoxia Response Element (HRE) RCGTG motif among observed DEGs in KS1 Patient iPSCs, KS1 Patient NSPCs, *Kmt2d^Δ/Δ^* HT22 cells, as well as KMT2D-bound genes in wild-type HT22 cells, and KMT2D-bound, down-regulated genes in *Kmt2d^Δ/Δ^* HT22 cells). Fisher’s Exact Test (*p<0.05, **p<0.01, ***p<0.001; ^†^FDR<0.05, ^††^FDR<0.01, ^†††^FDR<0.001).

**Supplementary Figure 5.**
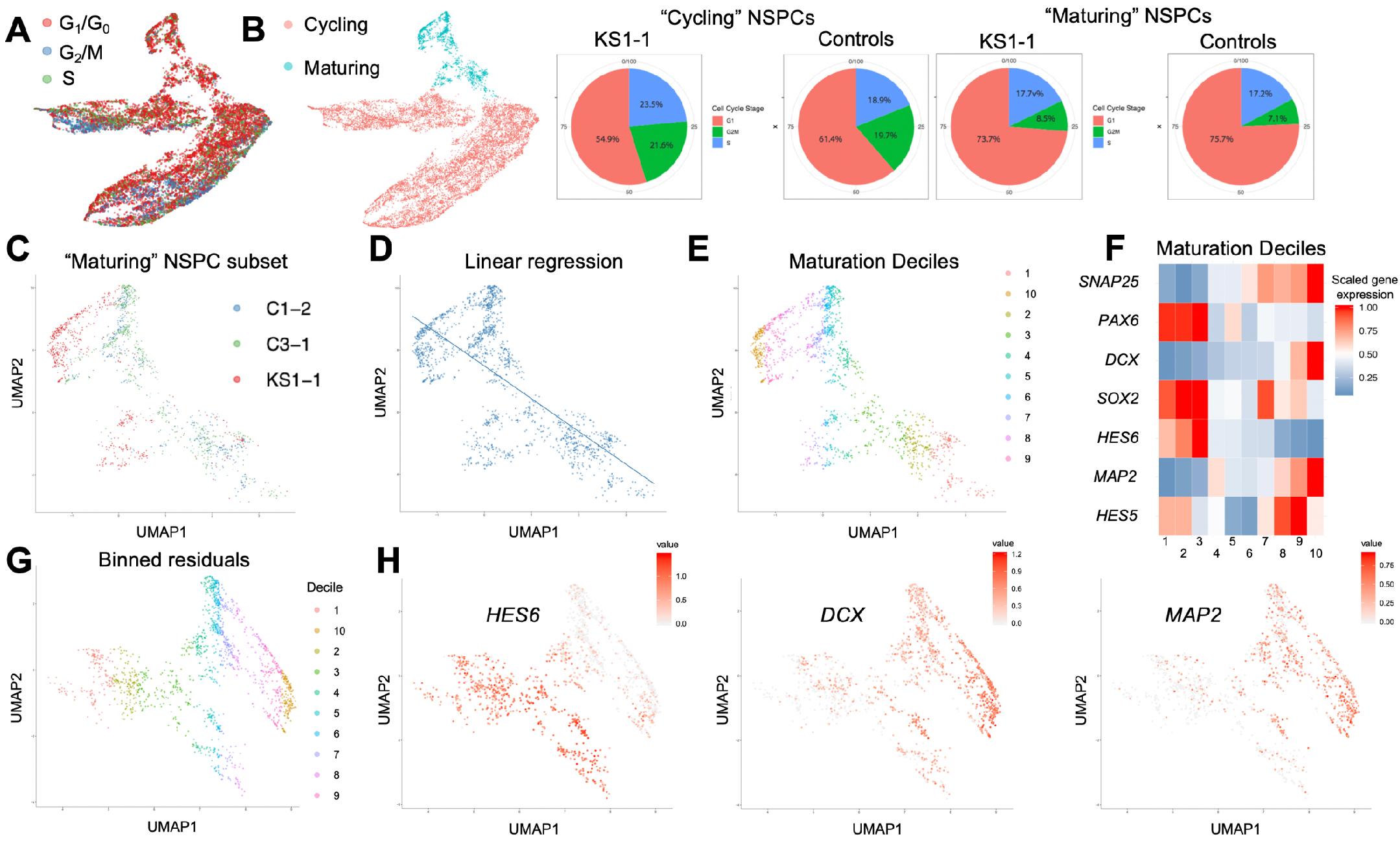
Stratified scRNA-seq analysis of NSPCs. Uniform Manifold Approximation Projection (UMAP) of single-cell NSPC libraries partitioned by (**A**) cell cycle marker expression into subsets of G_1_/G_0_, S, and G_2_/M cells, used for cycle phase-stratified differential expression analysis to rule out confounding differences in cell cycle phase composition on NSPC transcriptome comparisons. (**B**) Subset of “Cycling” versus non-cycling, “Maturing” NSPCs, which includes “Transitioning” and “Differentiating” cells as defined (Figure 4B), and UMAP-based cell cycle occupancies consistent with experimental FACS data (Figure 3F). (**C-H**) UMAP analysis of Differentiating NSPCs displaying (**C**) library patient ID’s, (**D**) smooth linear regression fitted to define the maturation trajectory and (**E**) binned deciles of progressively maturing cells along the regression. **(F)** Relative expression of selected NSPC markers defining directionality of the maturation trajectory. **(G)** Binned residuals used to calculate deciles containing equal number of cells along the axis of differentiation. (**H**) Representative NSPC marker expression plotted over binned residuals.

**Supplementary Figure 6.**
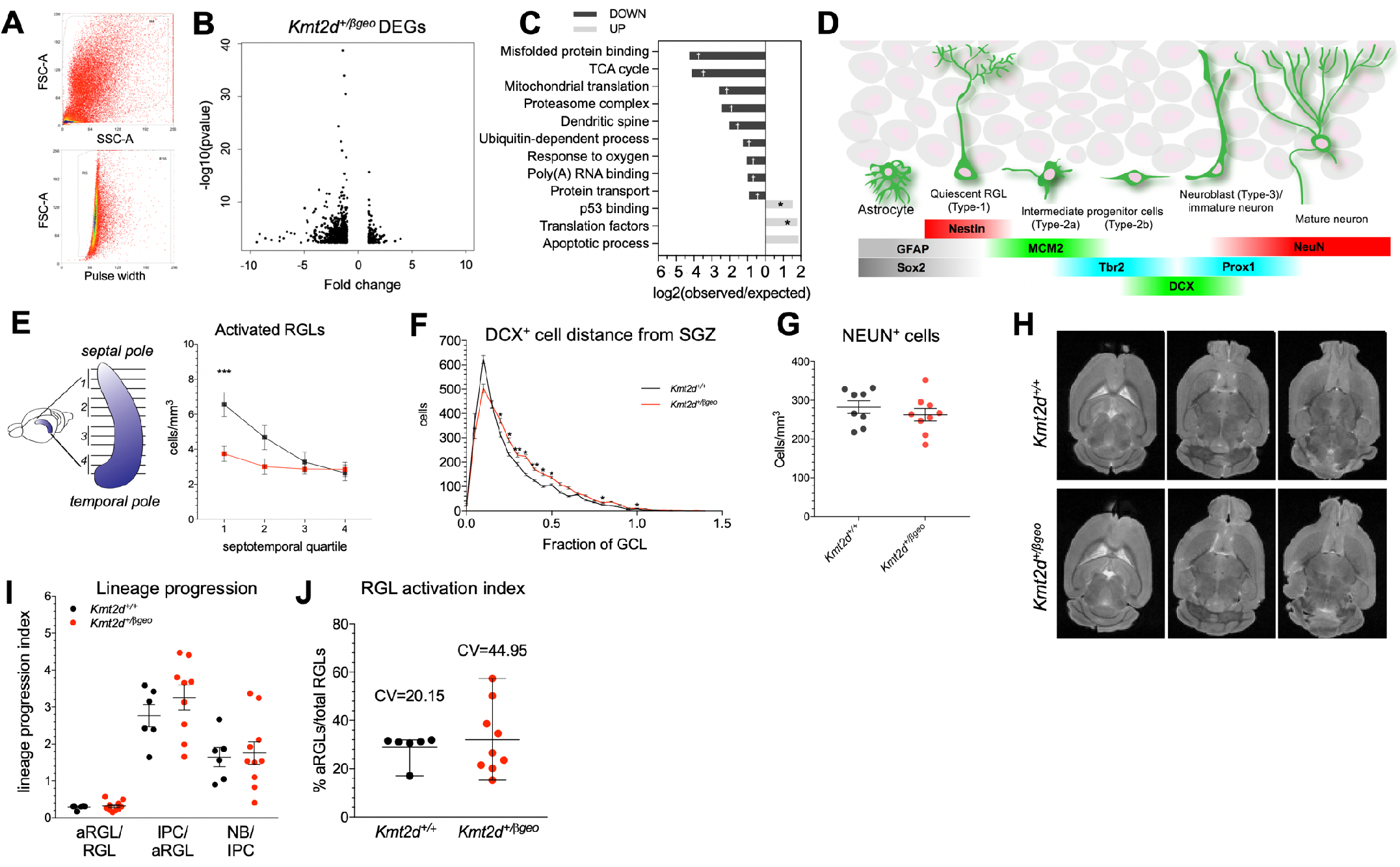
Phenotyping of *Kmt2d^+/βgeo^* mice. (**A**) Sample FACS gating for viable nuclei and doublet discrimination during purification of cycling EdU^+^ nuclei purified from *Kmt2d^+/+^* and *Kmt2d^+/βgeo^* mice at 16 hours post-EdU pulse for RNA-seq and cell cycle analysis. (**B**) RNA-seq analysis of differential gene expression in purified EdU^+^ DG nuclei from *Kmt2d^+/+^* and *Kmt2d^+/βgeo^* mice. (**C**) Gene networks most enriched among DEGs down- or up-regulated in *Kmt2d^+/βgeo^* nuclei, showing transcriptional suppression of cellular metabolic pathways. Fisher’s Exact Test (^†^FDR<0.05, ^††^FDR<0.01, ^†††^FDR<0.001). (**D**) Schematic depicting marker expression during sequential stages of adult DG neurogenesis. (**E**) Serial ordering of perfusion-fixed brain slices enables anatomically-stratified analysis of neurogenesis, for quantification of activated RGL NSPC density along the septotemporal axis of the DG in *Kmt2d^+/+^* and *Kmt2d^+/βgeo^* mice, indicating preferential disruption at the septal DG. Two-way ANOVA with post hoc multiple comparisons. (**F**) Quantification of DCX^+^ NB cell body distance from SGZ plane in 8-week-old mice (9-10 mice per genotype, >1,000 cells per mouse). Two-way ANOVA with post hoc multiple comparisons. (**G**) Quantification of RBFOX3/NEUN^+^ mature DG neurons in 8-week-old mice (8-9 mice per genotype, 10 z-stacks per mouse). Student’s t-test (n.s.). (**H**) Sample images of T2-weighted MRI (9.4T) in PFA-fixed brains of female mice 4 months old. (**I**) Comparison of lineage progression index, an approximation of expansion potential for each cell type transition, indicates absence of genotype-associated blockages at any particular cell-type transition analyzed, and (**J**) increased Coefficient of Variance (CV) in RGL activation rates in *Kmt2d^+/βgeo^* mice. Bars indicate mean ± SEM. (*p<0.05, **p<0.01, ***p<0.001).

**Supplementary Figure 7.**
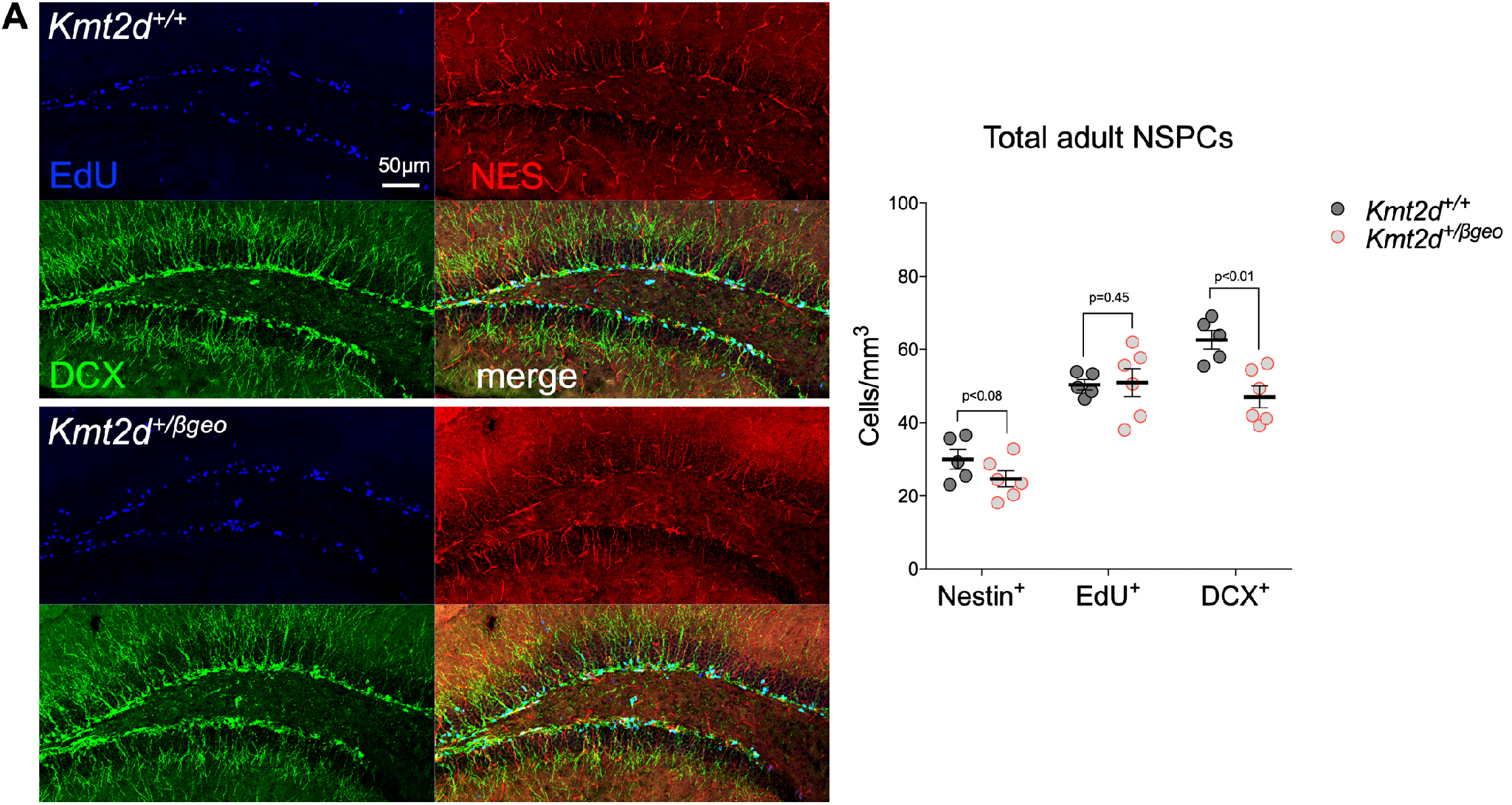
Pulse-labeling to birth-date adult-born NSPCs in vivo. (**A**) Representative immunostaining from *Kmt2d^+/+^* and *Kmt2d^+/βgeo^* mice (5-6 mice per genotype, 10 z-stacks per mouse) of EdU pulse-labeled cells extending a NES^+^ process (early RGL NSPCs) or DCX^+^ process (maturing NB NSPCs), showing the entire DG area quantified. Steady-state quantification of NSPCs and EdU-labeled NSPCs, confirming steady-state reduction of adult neurogenesis in *Kmt2d^+/βgeo^* mice, despite their increased number of EdU^+^DCX^+^ double-labeled NBs in the same experiment (Figure 5E-F). Bars indicate mean ± SEM, Student’s t-test. Scale bar 50 μm.

**Supplementary Figure 8.**
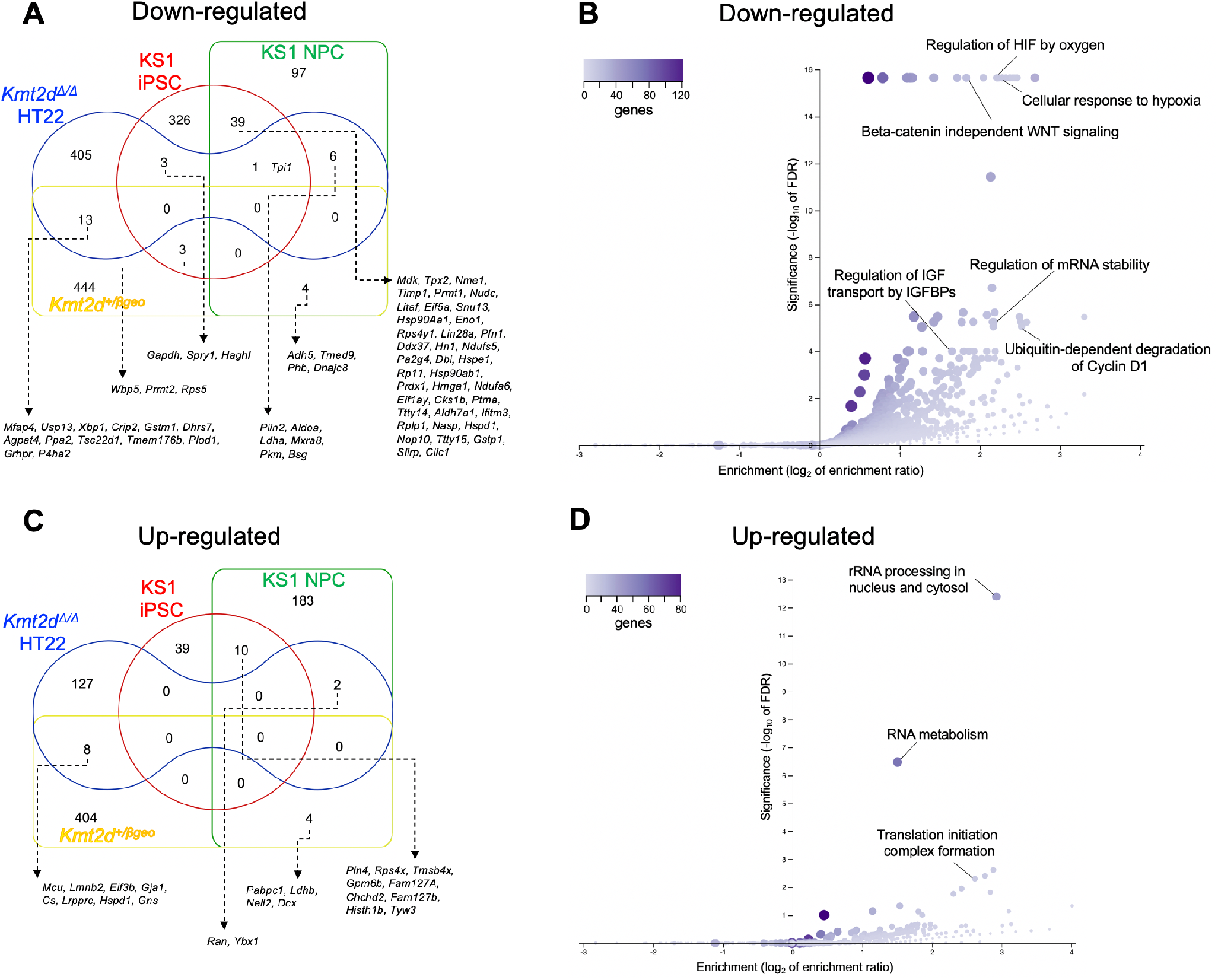
Comparison of gene expression across KS1 models. (**A-B**) Euler diagram depicting shared transcriptional downregulation in KS1 models with individual genes (**A**) and pathways enriched among down-regulated genes from all KS1 models presently studied (**B**). (**C-D**) Euler diagram depicting transcriptional upregulation in KS1 models with individual genes (**C**) and pathways enriched among up-regulated genes from all KS1 models presently studied (**D**). Enrichments expressed as log2 of enrichment ratio. Significance expressed as −log_10_ of FDR. (WebGestalt).

**Supplementary Figure 9.**
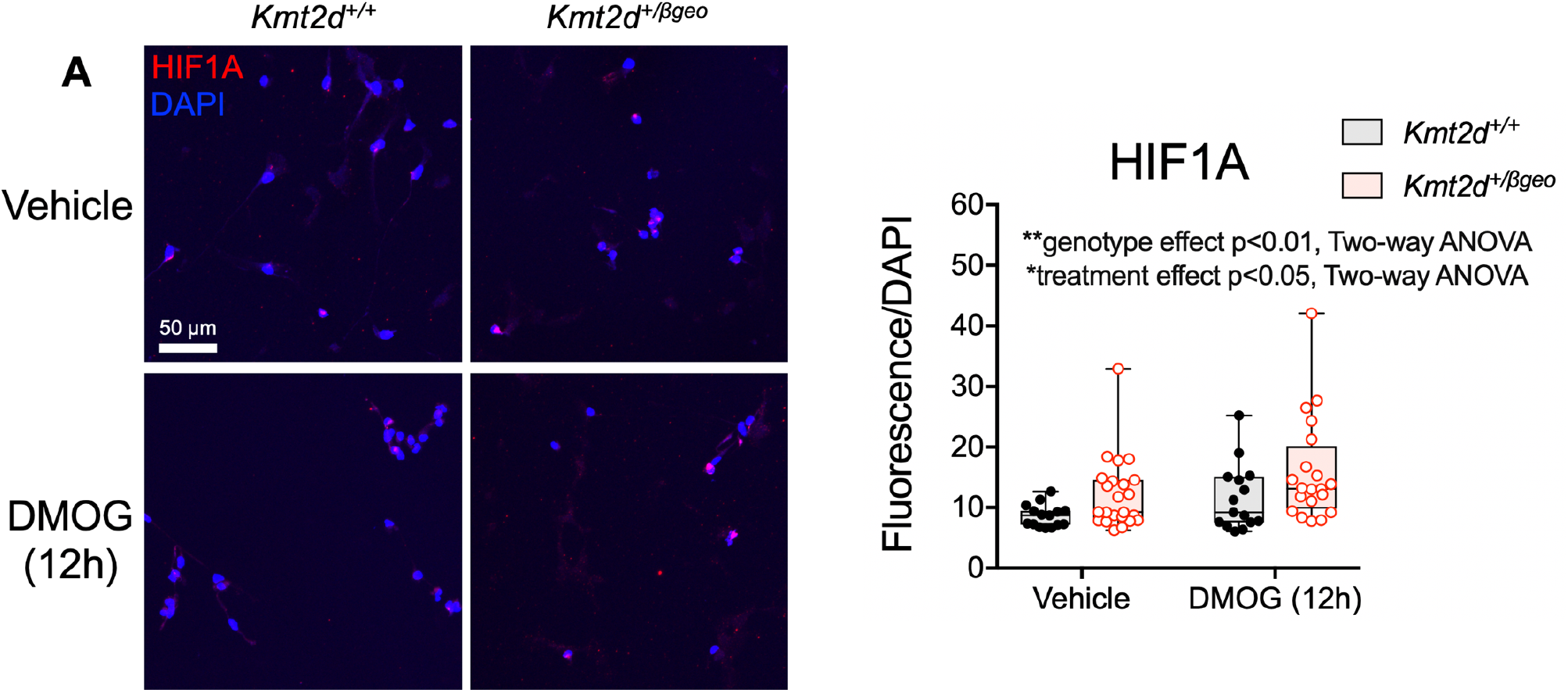
HIF1A activation in primary hippocampal NSPCs. (**A**) Representative confocal images of primary hippocampal NSPCs isolated from micro-dissected DG of *Kmt2d^+/+^* and *Kmt2d^+/βgeo^* mice, with quantification for analysis of HIF1A fluorescence inside the nucleus (DAPI^+^ volume). Two-way ANOVA with post-hoc multiple comparisons. (*p<0.05, **p<0.01, ***p<0.001). Boxes indicate mean ± interquartile range; whiskers indicate minima and maxima. Scale bar 50 μm.

**Supplementary Figure 10.**
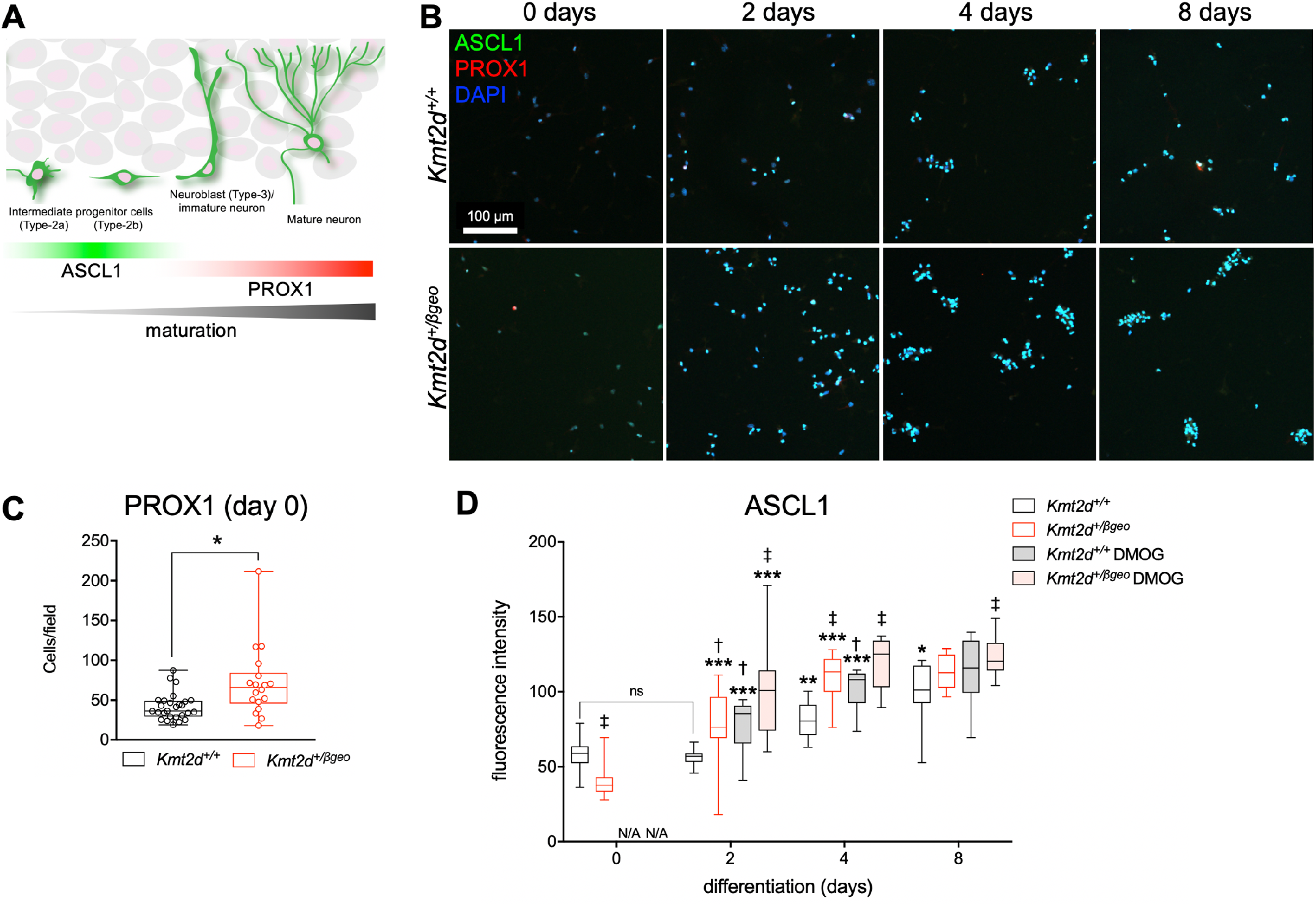
Precocious in vitro differentiation of primary hippocampal NSPCs. (**A**) Schematic depicting developmental expression of pro-neural transcription factor ASCL1 and maturing neuronal marker PROX1 in adult-born DG neurons. (**B**) Representative confocal images for analysis of NSPCs differentiating between 0 and 8 days in primary hippocampal NSPCs isolated from micro-dissected DG of *Kmt2d^+/+^* and *Kmt2d^+/βgeo^* mice, with quantifications (**C-D**). 22,307 cells analyzed individually across 176 fields of view. Two-way ANOVA with post hoc multiple comparisons. Boxes indicate mean ± interquartile range; whiskers indicate minima and maxima. (significance from previous time point *p<0.05, **p<0.01; ***p<0.001; significance from vehicle-treated wild-type ^†^p<0.05, ^‡^p<0.01). Scale bar 100 μm.

## Supplementary Tables

Supplementary Table 1: Differentially expressed genes in HT22 cells (*Kmt2d^+/+^* versus *Kmt2d^Δ/Δ^*) Supplementary Table 2: KMT2D ChIP-seq peaks in HT22 cells (*Kmt2d^+/+^* cells)

Supplementary Table 3: KMT2D-bound genes in HT22 cells (*Kmt2d^+/+^* cells) Supplementary Table 4: Differentially expressed genes in iPSCs (KS1 versus controls) Supplementary Table 5: Differentially expressed genes in NSPCs (KS1 versus controls) Supplementary Table 6: Intersected gene sets of iPSCs and NSPCs

Supplementary Table 7: Differentially expressed genes in NSPCs (stratified by subsets) Supplementary Table 8: Differentially expressed genes in EdU^+^ DG nuclei of *Kmt2d^+/βgeo^* mice Supplementary Table 9: MRI volumetric comparisons in wild-type and *Kmt2d^+/βgeo^* mice Supplementary Table 10: Reagents

## References

1. Schuettengruber B, Bourbon H-M, Di Croce L, Cavalli G. Genome regulation by polycomb and trithorax: 70 years and counting.. Cell 2017;171(1):34–57.

2. Bjornsson HT. The Mendelian disorders of the epigenetic machinery.. Genome Res. 2015;25(10):1473–1481.

3. Ng SB et al. Exome sequencing identifies MLL2 mutations as a cause of Kabuki syndrome.. Nat. Genet. 2010;42(9):790–793.

4. Miyake N et al. KDM 6 A Point Mutations Cause K abuki Syndrome.. Human mutation 2013;34(1):108–110.

5. Hannibal MC et al. Spectrum of MLL2 (ALR) mutations in 110 cases of Kabuki syndrome.. Am. J. Med. Genet. A 2011;155A(7):1511–1516.

6. Ang S-Y et al. KMT2D regulates specific programs in heart development via histone H3 lysine 4 di-methylation.. Development 2016;143(5):810–821.

7. Ortega-Molina A et al. The histone lysine methyltransferase KMT2D sustains a gene expression program that represses B cell lymphoma development.. Nat. Med. 2015;21(10):1199–1208.

8. Zhang J et al. Disruption of KMT2D perturbs germinal center B cell development and promotes lymphomagenesis.. Nat. Med. 2015;21(10):1190–1198.

9. Lee J-E et al. H3K4 mono- and di-methyltransferase MLL4 is required for enhancer activation during cell differentiation.. Elife 2013;2:e01503.

10. Lin-Shiao E et al. KMT2D regulates p63 target enhancers to coordinate epithelial homeostasis.. Genes Dev. 2018;32(2):181–193.

11. Froimchuk E, Jang Y, Ge K. Histone H3 lysine 4 methyltransferase KMT2D.. Gene 2017;627:337– 342.

12. Bjornsson HT et al. Histone deacetylase inhibition rescues structural and functional brain deficits in a mouse model of Kabuki syndrome.. Sci Transl Med 2014;6(256):256ra135.

13. Boisgontier J et al. Anatomical and functional abnormalities on MRI in kabuki syndrome.. Neuroimage Clin. [published online ahead of print: November 19, 2018]; doi:10.1016/j.nicl.2018.11.020

14. Gonçalves JT, Schafer ST, Gage FH. Adult neurogenesis in the hippocampus: from stem cells to behavior.. Cell 2016;167(4):897–914.

15. Moreno-Jiménez EP et al. Adult hippocampal neurogenesis is abundant in neurologically healthy subjects and drops sharply in patients with Alzheimer’s disease.. Nat. Med. 2019;25(4):554–560.

16. Tobin MK et al. Human hippocampal neurogenesis persists in aged adults and alzheimer’s disease patients.. Cell Stem Cell 2019;24(6):974–982.e3.

17. Sorrells SF et al. Human hippocampal neurogenesis drops sharply in children to undetectable levels in adults.. Nature 2018;555(7696):377–381.

18. Berg DA et al. A common embryonic origin of stem cells drives developmental and adult neurogenesis.. Cell 2019;177(3):654–668.e15.

19. Zheng X et al. Metabolic reprogramming during neuronal differentiation from aerobic glycolysis to neuronal oxidative phosphorylation.. Elife 2016;5. doi:10.7554/eLife.13374

20. Mazumdar J et al. O_2_ regulates stem cells through Wnt/β-catenin signalling.. Nat. Cell Biol. 2010;12(10):1007–1013.

21. Chatzi C, Schnell E, Westbrook GL. Localized hypoxia within the subgranular zone determines the early survival of newborn hippocampal granule cells.. Elife 2015;4:e08722.

22. Morimoto BH, Koshland DE. Induction and expression of long- and short-term neurosecretory potentiation in a neural cell line.. Neuron 1990;5(6):875–880.

23. Wang C et al. Enhancer priming by H3K4 methyltransferase MLL4 controls cell fate transition.. Proc. Natl. Acad. Sci. USA 2016;113(42):11871–11876.

24. Guo C et al. Global identification of MLL2-targeted loci reveals MLL2’s role in diverse signaling pathways.. Proc. Natl. Acad. Sci. USA 2012;109(43):17603–17608.

25. Semenza GL. HIF-1: mediator of physiological and pathophysiological responses to hypoxia.. J. Appl. Physiol. 2000;88(4):1474–1480.

26. Guimarães-Camboa N et al. Hif1α represses cell stress pathways to allow proliferation of hypoxic fetal cardiomyocytes.. Dev. Cell 2015;33(5):507–521.

27. Benita Y et al. An integrative genomics approach identifies Hypoxia Inducible Factor-1 (HIF-1)-target genes that form the core response to hypoxia.. Nucleic Acids Res. 2009;37(14):4587–4602.

28. Sanz JH, Lipkin P, Rosenbaum K, Mahone EM. Developmental profile and trajectory of neuropsychological skills in a child with Kabuki syndrome: implications for assessment of syndromes associated with intellectual disability.. Clin Neuropsychol 2010;24(7):1181–1192.

29. Wen Z et al. Synaptic dysregulation in a human iPS cell model of mental disorders.. Nature 2014;515(7527):414–418.

30. Li W et al. Rapid induction and long-term self-renewal of primitive neural precursors from human embryonic stem cells by small molecule inhibitors.. Proc. Natl. Acad. Sci. USA 2011;108(20):8299– 8304.

31. McInnes L, Healy J, Saul N, Großberger L. UMAP: uniform manifold approximation and projection. JOSS 2018;3(29):861.

32. Benjamin JS et al. A ketogenic diet rescues hippocampal memory defects in a mouse model of Kabuki syndrome.. Proc. Natl. Acad. Sci. USA 2017;114(1):125–130.

33. Morris AM, Churchwell JC, Kesner RP, Gilbert PE. Selective lesions of the dentate gyrus produce disruptions in place learning for adjacent spatial locations.. Neurobiol. Learn. Mem. 2012;97(3):326– 331.

34. Zhuo J-M et al. Young adult born neurons enhance hippocampal dependent performance via influences on bilateral networks.. Elife 2016;5. doi:10.7554/eLife.22429

35. Snyder JS. Recalibrating the relevance of adult neurogenesis.. Trends Neurosci. 2019;42(3):164– 178.

36. Chakraborty AA et al. Histone demethylase KDM6A directly senses oxygen to control chromatin and cell fate.. Science (80-.). 2019;363(6432):1217–1222.

37. Batie M et al. Hypoxia induces rapid changes to histone methylation and reprograms chromatin.. Science (80-.). 2019;363(6432):1222–1226.

38. Luo W, Wang Y. Epigenetic regulators: multifunctional proteins modulating hypoxia-inducible factor-α protein stability and activity.. Cell Mol. Life Sci. 2018;75(6):1043–1056.

39. Bernas S, Leiter O, Walker T, Kempermann G. Isolation, culture and differentiation of adult hippocampal precursor cells. Bio Protoc 2017;7(21). doi:10.21769/BioProtoc.2603

40. Patro R, Duggal G, Love MI, Irizarry RA, Kingsford C. Salmon provides fast and bias-aware quantification of transcript expression.. Nat. Methods 2017;14(4):417–419.

41. Soneson C, Love MI, Robinson MD. Differential analyses for RNA-seq: transcript-level estimates improve gene-level inferences. [version 2; peer review: 2 approved]. F1000Res. 2015;4:1521.

42. Robinson MD, McCarthy DJ, Smyth GK. edgeR: a Bioconductor package for differential expression analysis of digital gene expression data.. Bioinformatics 2010;26(1):139–140.

43. McCarthy DJ, Chen Y, Smyth GK. Differential expression analysis of multifactor RNA-Seq experiments with respect to biological variation.. Nucleic Acids Res. 2012;40(10):4288–4297.

44. Ritchie ME et al. limma powers differential expression analyses for RNA-sequencing and microarray studies.. Nucleic Acids Res. 2015;43(7):e47.

45. Giner G, Smyth GK. Statmod: probability calculations for the inverse Gaussian distribution.. arXiv preprint arXiv:1603.06687 2016;

46. Love MI, Huber W, Anders S. Moderated estimation of fold change and dispersion for RNA-seq data with DESeq2.. Genome Biol. 2014;15(12):550.

47. Trapnell C et al. The dynamics and regulators of cell fate decisions are revealed by pseudotemporal ordering of single cells.. Nat. Biotechnol. 2014;32(4):381–386.

48. Lun ATL, McCarthy DJ, Marioni JC. A step-by-step workflow for low-level analysis of single-cell RNA-seq data with Bioconductor. [version 2; peer review: 3 approved, 2 approved with reservations]. F1000Res. 2016;5:2122.

49. Langmead B, Salzberg SL. Fast gapped-read alignment with Bowtie 2.. Nat. Methods 2012;9(4):357–359.

50. Zhang Y et al. Model-based analysis of ChIP-Seq (MACS).. Genome Biol. 2008;9(9):R137.

51. Amemiya HM, Kundaje A, Boyle AP. The ENCODE blacklist: identification of problematic regions of the genome.. Sci. Rep. 2019;9(1):9354.

52. Liao Y, Smyth GK, Shi W. The R package Rsubread is easier, faster, cheaper and better for alignment and quantification of RNA sequencing reads.. Nucleic Acids Res. 2019;47(8):e47.

53. Landt SG et al. ChIP-seq guidelines and practices of the ENCODE and modENCODE consortia.. Genome Res. 2012;22(9):1813–1831.

54. Rainer. EnsDb.Mmusculus.v79: Ensembl based annotation package.. R package version 2.99.0. 2017;

55. Lawrence M et al. Software for computing and annotating genomic ranges.. PLoS Comput. Biol. 2013 9(8):e1003118.

56. Rizzardi LF et al. Neuronal brain-region-specific DNA methylation and chromatin accessibility are associated with neuropsychiatric trait heritability.. Nat. Neurosci. 2019;22(2):307–316.

57. Picelli S et al. Smart-seq2 for sensitive full-length transcriptome profiling in single cells.. Nat. Methods 2013;10(11):1096–1098.

58. Kim D, Langmead B, Salzberg SL. HISAT: a fast spliced aligner with low memory requirements.. Nat. Methods 2015;12(4):357–360.

59. Mudge JM, Harrow J. Creating reference gene annotation for the mouse C57BL6/J genome assembly.. Mamm. Genome 2015;26(9–10):366–378.

60. Trapnell C et al. Differential gene and transcript expression analysis of RNA-seq experiments with TopHat and Cufflinks.. Nat. Protoc. 2012;7(3):562–578.

61. Wang J, Vasaikar S, Shi Z, Greer M, Zhang B. WebGestalt 2017: a more comprehensive, powerful, flexible and interactive gene set enrichment analysis toolkit.. Nucleic Acids Res. 2017;45(W1):W130–W137.

